# Revealing dynamic temporal trajectories and underlying regulatory networks with *Cflows*

**DOI:** 10.1101/2023.03.28.534644

**Authors:** Xingzhi Sun, Shabarni Gupta, Alexander Tong, Manik Kuchroo, Dhananjay Bhaskar, Chen Liu, Aarthi Venkat, Beatriz P. San Juan, Laura Rangel, Vanina Rodriguez, John G. Lock, Christine L. Chaffer, Smita Krishnaswamy

## Abstract

While single-cell technologies provide snapshots of tumor states, building continuous trajectories and uncovering causative gene regulatory networks remains a significant challenge. We present *Cflows*, an AI framework that combines neural ODE networks with Granger causality to infer continuous cell state transitions and gene regulatory interactions from static scRNA-seq data. In a new 5-time point dataset capturing tumorsphere development over 30 days, *Cflows* reconstructs two types of trajectories leading to tumorsphere formation or apoptosis. Trajectory-based cell-of-origin analysis delineated a novel cancer stem cell profile characterized by CD44^*hi*^EPCAM^+^CAV1^+^, and uncovered a cell cycle–dependent enrichment of tumorsphere-initiating potential in G2/M or S-phase cells. *Cflows* uncovers ESRRA as a crucial causal driver of the tumor-forming gene regulatory network. Indeed, ESRRA inhibition significantly reduces tumor growth and metastasis *in vivo. Cflows* offers a powerful framework for uncovering cellular transitions and dynamic regulatory networks from static single-cell data.

## Introduction

Cancer is a highly dynamic biological system. Although single-cell technologies have provided valuable snapshots of gene expression within tumors at different stages, these snapshots are static and fail to capture the continuously evolving transcriptional networks over time (Figure 1a). Moreover, computational approaches to seamlessly connect evolving cell states into pathogenic trajectories and define their underlying causative gene regulatory networks remain underdeveloped. Consequently, very little is known about the molecular programs of the specific cells that initiate tumors or the dynamic transcriptional programs that enable them to drive cancer progression. This knowledge is essential for predicting causal molecular programs that drive disease progression, and ultimately, for defining new therapeutic strategies to stop it.

**Figure 1:**
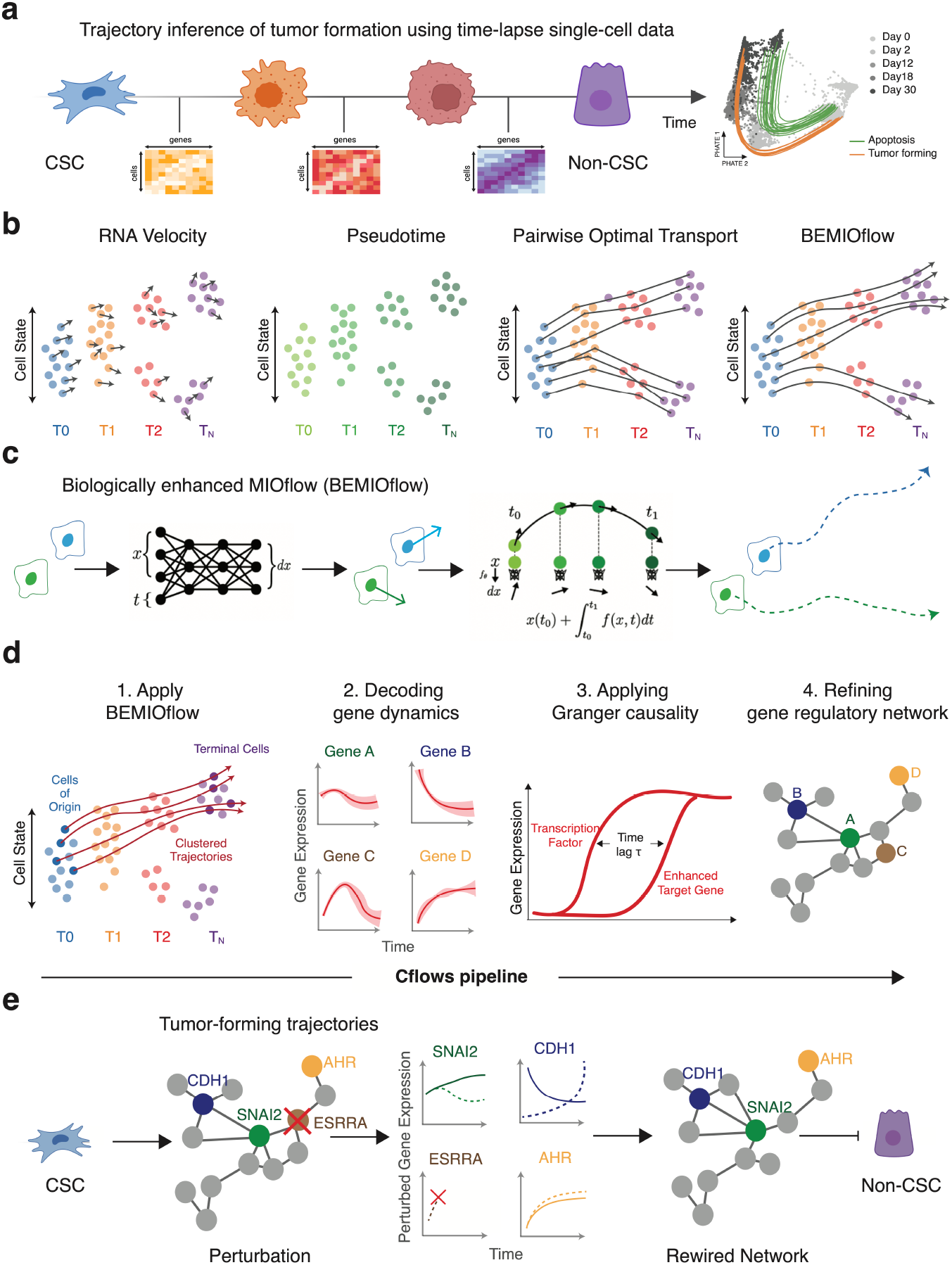
Overview of the Cflows framework. **a**, Time-lapse single-cell RNA-seq of tumorsphere formation. BEMIOflow, the trajectory-inference module of the Cflows pipeline, recovers two groups of continuous trajectories: tumor-forming and apoptotic. **b,** Existing methods such as RNA Velocity, pseudotime estimation, and pairwise optimal transport provide coarse or population-level representations of dynamic cellular processes. In contrast, BEMIOflow infers continuous, single-cell trajectories over time. **c**, BEMIOflow models cell trajectories using neural ordinary differential equations (Neural ODEs). At each time point, a neural network estimates the instantaneous derivative of each cell’s state, which is then integrated over time to generate continuous trajectories. **d**, Overview of the Cflows pipeline. 1) BEMIOflow infers cell trajectories using a novel proliferation-rate-aware architecture. 2) Cell trajectories are decoded to gene dynamics. 3) Granger-causality analysis (GC) derives gene-regulatory networks. 4) Predicted edges are cross-referenced with public databases to increase confidence. **e**, GC-derived regulatory network highlights ESRRA as a central transcription factor controlling key tumor-formation genes (including CDH1, SNAI1, AHR). ESSRA is required to maintain the cancer stem cell (CSC) state, and in vivo tumor formation and metastasis.

Current methods, including RNA velocity [1] and pseudotime [2], attempt to imbue single-cell analysis with dynamic information. RNA velocity predicts a velocity arrow of a cell based on the ratio of unspliced to spliced mRNA, which is often very noisy, while pseudotime orders all cells along a single trajectory based on manifold structure. However, these methods do not infer continuous trajectories at the single-cell level across the cellular data (Figure 1b). In this work, we show that deep-learning-based flow models [3–6] can be crucial in developing an analysis pipeline that subsumes RNA-velocity and pseudotime and predicts a flow, or continuous-time transitions, of individual cells from temporally sampled data.

We developed *Cflows*, or **Causality-from-flows**, a pipeline that combines a neural flow-based model of dynamic optimal transport called *BEMIOflow*, or **Biology-Enhanced MIOflow** along with causality and analysis of gene networks (Figure 1b-d). Our previous work, *MIOflow* [4], showcased the use of neural ordinary differential equations (ODEs) [7, 8] to infer continuous dynamics from static snapshot single-cell data. Neural ODEs, unique in their ability to produce time-varying derivatives, represent changes in cell state or position over time, ensuring accurate cellular trajectory inference. The *Cflows* pipeline starts with *BEMIOflow* (Figure 1c), which enhances our previous work *MIOflow* that facilitated dynamic and optimal cell transport from one time point to another. However, BEMIOflow incorporates another neural network for predicting the growth rate of a cell continuously in time to model cell growth and death to augment the optimal transport. *Cflows* further integrates these neural ODE dynamics with time-lapsed causality analysis [9] via granger causality to construct detailed gene-regulatory networks and identify potential therapeutic targets. Unlike traditional network-building paradigms that rely on co-expression as a proxy for association, *Cflows* captures the time lag in causal regulatory interactions, enabling the discovery of evolving gene regulatory networks (Figure 1d).

To demonstrate *Cflows*’s ability to uncover continuous single-cell time trajectories governing tumor initiation and progression, as well as the underlying causal gene regulatory networks, we first designed a series of detailed benchmarking experiments. We simulated data using SERGIO [10] with known ground truth, then compared *Cflows* to other methods for trajectory prediction and network inference, showing that it vastly outperforms existing methods.

Next we applied *Cflows* to study tumor initiation and progression using triple-negative breast cancer (TNBC) as a model system. Cancer stem cells (CSCs) are cancer cells uniquely endowed with the ability to initiate tumors, however, the marker profiles to enrich for these specialized cells are poorly defined, as are the molecular programs and temporal trajectories that enable a single CSC to generate a robustly growing tumor. To study this, we seeded single cells from a heterogeneous TNBC cell line containing CSCs into a 3D culture and measured tumorsphere initiation and growth by single cell RNA-sequencing at five timepoints spanning 30 days. From this data, Cflows infers plausible trajectories from initiation to subsequent stages of tumorsphere growth. Accordingly, we present a new detailed understanding of CSC biology, including a refined cell surface marker (CD44+EPCAM+CAV1+) profile and cell cycle profile (G2/M or S-phase enriched) of CSCs that initiate tumorspheres. Moreover, by following pathogenic CSC trajectories forward, we reveal the first causative gene regulatory network that drives tumor progression, of which a core molecular component is regulated by estrogen receptor related alpha (ESRRA). Indeed, we show that genetic ablation, or drugs that target ESRRA, disrupt tumorsphere formation across an array of TNBC cell lines. Moreover, we show that inhibiting ESRRA significantly inhibits primary tumor growth and metastasis *in vivo*. Targeting ESRRA warrants further investigation as a new therapeutic strategy to prevent metastatic growth in TNBC.

*Cflows* is a state-of-the-art pipeline for learning continuous cellular trajectories and decoding causal gene regulatory networks. This study represents the first experimentally validated study of a flow-based trajectory inference model for biological dynamics predictions including novel target finding. In this TNBC system we derive two types of trajectories, and further demonstrate how such trajectories can be used to derive gene regulatory networks whose modules present new potential targets for treatment.

## Results

### Overview of the Cflows pipeline for learning cellular dynamics and underlying causal gene regulatory networks

Time-lapse single-cell data (Figure 1a) poses significant computational challenges. In cancer, for example, cells undergo significant and rapid transcriptional changes in response to natural tumor evolution. Single cell data measures distributions of cells at discrete timepoints that largely do not overlap in cellular state with previous or subsequent timepoints. Under these circumstances, previously presented techniques for inferring dynamics such as RNA velocity [1] and pseudotime [2] are unable to infer cellular evolution at a single cell scale as these approaches rely on well-sampled manifolds without large gaps in cell state space or time. Most critically, however, these approaches do not help users understand the gene regulatory dynamics necessary to drive these trajectories forward. Extracting longitudinal dynamics at the single cell level, and transcriptional drivers leading to heterogeneous cell fates, can be challenging as there are no methods that can interpolate between irregularly spaced data at a single cell level and learn gene-gene regulatory dynamics. In our previous work, we pioneered learning cellular dynamics from disconnected snapshot single-cell data via neural ODE networks [3, 4] (Figure 1b). However, these methods have not been used for intensive gene network discovery in a biological setting. Here we developed Cflows, or Causality-from-flows, that learns cellular and gene trajectories from disconnected single-cell data (Figure 1d), and then exploits these to identify the *transcriptional networks* (Figure 1e) underlying cellular dynamics.

The Cflows pipeline consists of the following steps:

1. Biologically Enhanced MIOflow (BEMIOflow) for learning cellular trajectories
2. Decoding transcriptional programs and gene dynamics
3. Gene regulatory network inference using Granger Causality
4. Integrating established databases to increase trust in gene regulatory relationships

### 1. Biologically Enhanced MIOflow for learning cellular trajectories

Inferring time-varying cellular dynamics from scRNA-seq data presents two fundamental challenges. First, because the measurements are destructive, each cell is profiled only once, precluding longitudinal tracking and leaving no ground-truth correspondences across time points. Second, measurements are obtained at discrete intervals, limiting our ability to infer continuous trajectories.

Several methods have been proposed to address these limitations, but each has key shortcomings. *Optimal transport* [11] methods infer pairings but lack a continuous model of temporal evolution. *RNA velocity* [1] estimates local velocity vectors but does not provide global trajectories and often yields inconsistent directions over neighboring times. *Pseudotime* algorithms [2] provide coarse-grained developmental orderings but cannot infer individual cell trajectories.

To overcome these limitations, we previously developed MIOflow [4], which combines dynamic optimal transport with neural ODEs [7]. By minimizing transport cost between distributions at successive time points, MIOflow infers smooth trajectories between cellular populations such that one population flows to the next and matches timepoints. The learned neural ODE can then generate smooth trajectories for individual cells over time, starting from the initial timepoint. Although MIOflow addresses both of the aforementioned challenges, it assumes a fixed population size and does not account for cell proliferation or death, which are fundamental to many biological systems, especially cancer. Furthermore, in order for the cellular trajectories to be plausible, it is important that they adhere to the cellular manifold. While MIOflow attempts to address this by utilizing a multi-scale diffusion latent space, we hypothesize that the PHATE dimensionality reduction method is more effective at learning low dimensional manifolds.

Therefore, we introduce Biologically Enhanced MIOflow (BEMIOflow, Figure 1c), which extends MIOflow in two critical ways and forms the first stage of the *Cflows* pipeline. First, BEMIOflow operates in PHATE [12] space, a low-dimensional embedding space that preserves manifold geometry and emphasizes developmental continuity. Second, it explicitly models cellular proliferation and death, allowing for realistic, non-conservative dynamics, and enables a novel cell-of-origin analysis. These enhancements, summarized below, enable BEMIOflow to infer single-cell trajectories that more faithfully reflect the underlying biology and capture complex temporal dynamics. We illustrate BEMIOflow with a schematic in Figure S1.

- **BEMIOflow infers trajectory on PHATE space.** First, we perform trajectory training in PHATE [12] space – a dimensionality-reduced representation that preserves the intrinsic structure, geometry, and distances of the gene expression manifold. This embedding enables the accurate identification of cellular progression paths while quantifying energetic costs, ultimately leading to the inference of trajectories that are both energy-efficient and biologically plausible. To embed the gene expression data into the PHATE space, we first reduced its dimensionality with Principal Component Analysis (PCA) and then trained an encoder neural network regularized by PHATE distances [13].
- **BEMIOflow enhances temporal resolution of data for more accurate trajectory inference.** Second, we improve the temporal resolution by subclustering cells at each time point, thereby creating refined pseudotime bins that capture subtle progression dynamics. This finer-grained temporal binning enables more accurate trajectory inference than relying solely on experimental time points.
- **BEMIOflow Models Growth/Death Rate And Enables Cell-Of-Origin Analysis.** In biological systems, a cell can proliferate into multiple cells or undergo cell death – processes that are particularly significant in cancer. This necessitates a model capable of accurately capturing growth and death along each trajectory, rather than assuming that each cell simply transitions state. Furthermore, in cancer research, it is critical to determine which early cell states are capable of tumor-initiation, an analysis we refer to as **cell-of-origin analysis**. In BEMIOflow, we incorporate an auxiliary network that estimates cellular growth/death rates to reflect realistic dynamics and differentiation. These rates help the model to account for changes in cell numbers along the trajectories, ensuring a better match with the observed data. Furthermore, we initialize trajectories from cell states exhibiting high growth rates. After training, we track the trajectories that culminate in states of interest (e.g., tumor-forming or apoptotic) back to their starting points, thereby revealing their origins.

### 2. Decoding gene dynamics and transcriptional programs

To translate the low-dimensional cell state trajectories inferred by BEMIOflow into functionally informative gene dynamics, we projected them back to the original gene space. This was accomplished using a neural network decoder followed by an inverse PCA decoder, yielding dynamic expression profiles for each gene at the single-cell level (Figure S1).

Furthermore, to provide an overall understanding of the dynamic transcriptional programs, we systematically ordered and grouped genes by their dynamic expression patterns: First, we captured the aggregate dynamics by computing the mean expression for each gene across all cells over time and rescaling the values to a range of [0, 1]. We term this normalized profile a gene trend. Next, we identified the point of maximum expression for each trend, or its peak time, to pinpoint when a gene is most active. Ordering the genes by their peak times revealed a temporal cascade of gene activation, from early-to late-peaking. Finally, we applied kernel density estimation to the distribution of peak times to partition genes based on peak-time. The modes of the resulting density, separated by local minima, correspond to distinct groups of co-activated genes, which we define as gene clusters.

### 3. Gene Regulatory Network Inference using Granger Causality

After obtaining the clustered gene trends, Cflows infers the underlying causal gene regulatory networks along each cellular trajectory using a time-lagged inference method called Granger causality (Figure 1d). Although many existing methods, such as those based on correlation or mutual information [14], assume that cellular states are static, we contend that the incorporation of dynamic cellular information is essential to uncover the true transcriptional network. Because BEMIOflow outputs complete transcriptional trends for each gene over time, Cflows can infer more accurate gene-gene relationships using Granger causality. Unlike simple correlation measures, Granger causality tests whether past values of a transcription factor *X* improve the prediction of gene *Y*, thereby revealing directional regulatory influences. Based on this directional forecasting, we further determine whether transcription factor *X* enhances or represses gene *Y*, a measure we term the **Total Granger Causality Score** (TGCS) (see Methods).

To enhance the biological plausibility of the inferred gene regulatory relationships, we impose two key constraints. First, we restrict potential interactions by considering only highly variable transcription factors as regulators and highly variable genes as targets. Second, we enforce a temporal constraint by limiting regulatory interactions to cases where the source (regulatory) clusters peak earlier than the target (regulated) clusters.

### Integrating established databases to increase trust in gene regulatory relationships

Lastly, *Cflows* enriches the inferred gene regulatory network by cross-linking the interactions uncovered by the TGCS with established gene regulation databases (Figure 1e). This integration produces a subnetwork of highly plausible transcription factor-gene relationships, which we then leverage to elucidate the causative networks that drive tumorsphere formation and progression, or that to lead to apoptosis. We then modulate key components of the gene regulatory networks to confirm causality. Here, we identify key drivers of tumor-initiation and validate causality in *in vitro* models of tumorsphere formation and in *in vivo* models of primary tumor growth and metastasis.

For a more in-depth explanation of *Cflows* and it’s improved implementation, Granger-causality gene network inference and all comparisons on synthetic data please see Methods.

#### *Cflows* accurately infers trajectories and causal transcriptional networks that outperform existing methods

Through rigorous comparisons, we show that *Cflows* not only learns cellular trajectories better than other techniques, but also more accurately infers known gene regulatory relationships than established methods that do not incorporate cellular dynamics (Figure 2).

**Figure 2:**
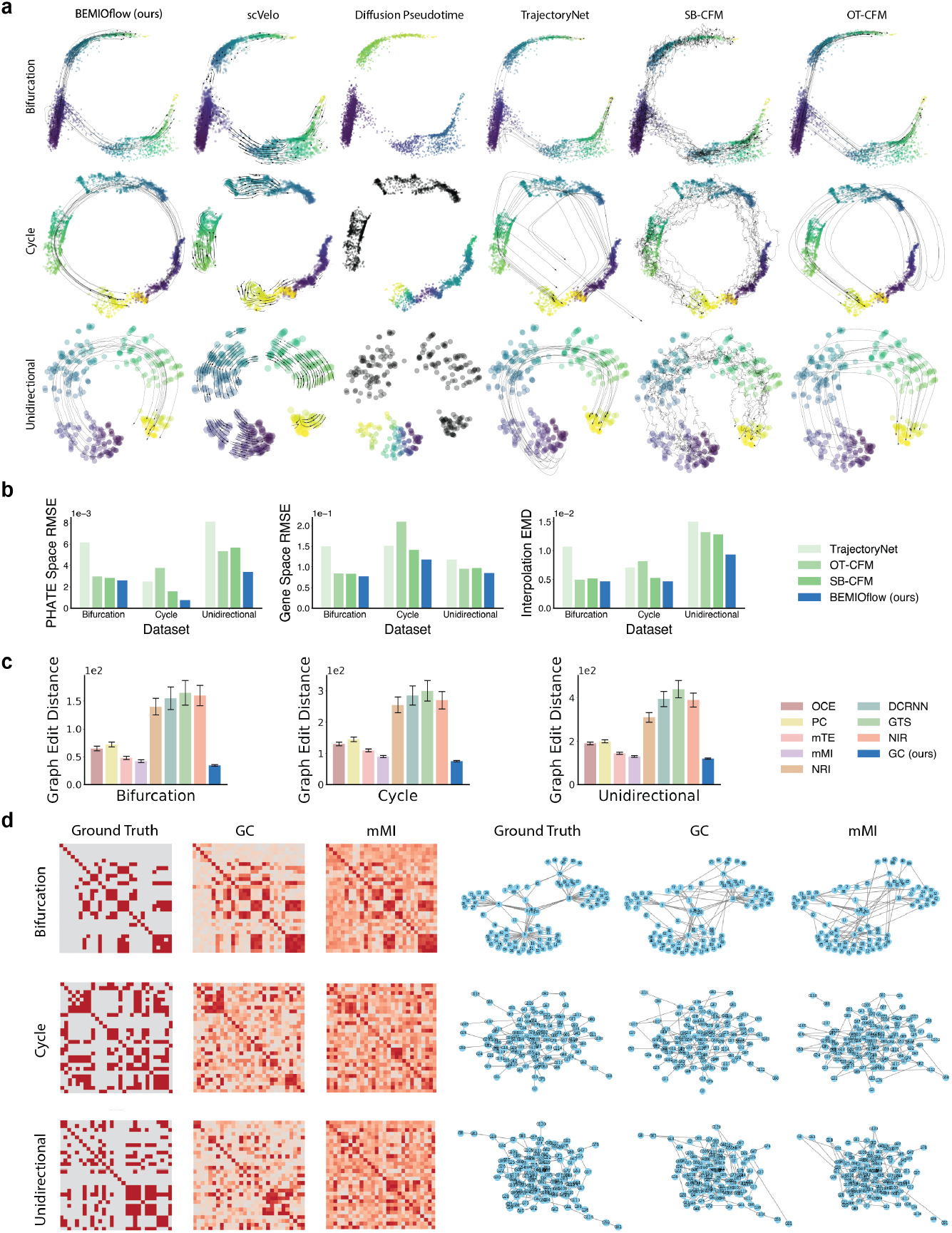
Benchmarking *Cflows* for trajectory and regulatory-network inference. **a**, Single-cell trajectory snapshots simulated with SERGIO (ground truth) and trajectories inferred by six algorithms. BEMIOflow, the trajectory-inference module of *Cflows*, recovers continuous, manifold-conforming paths for individual cells. By contrast, scVelo provides only local and oftentimes contradictory velocity vectors, Diffusion Pseudotime collapses all cells onto a single path and omits disconnected populations, and TrajectoryNet, SB-CFM and OT-CFM yield paths that deviate from the cellular manifold. **b**, Quantitative assessment of trajectory accuracy. Left, root mean-squared error (RMSE) between inferred and true coordinates in PHATE space. Center, RMSE in the original gene-expression space. Right, earth-mover’s distance (EMD) between predicted and true expression distributions at a held-out intermediate time point. Lower values indicate better performance; BEMIOflow attains the lowest error across all metrics. **c**, Mean graph-edit distance (GED) between inferred and ground-truth networks of increasing size. GC consistently outperforms OCE, PC, mTE, nMI, NRI, DCRNN, GTS and NIR (mean ± s.d. computed using n = 5 sets of trajectories). **d**, Ground-truth regulatory network simulated using SERGIO (left) and networks inferred by the methods. Granger causality (GC), the network-inference module of Cflows, most closely matches the true graph.

We used the SERGIO [10] simulator (see Figure 2) to generate three ground truth transcriptional networks and gene expression data describing three scenarios: bifurcation, unidirectional, and cyclical. We began by creating a reaction network between genes, which was subsequently simulated using stochastic dynamics. Single-cell data were created by sampling cells from gene expression over time.

To add realism, technical noise was introduced to the gene counts. The resulting data matrices served as inputs for Cflows and other analytical algorithms.

We compared BEMIOflow, part of the Cflows pipeline, to TrajectoryNet [3] and recent flow matching based methods such as optimal transport conditional flow matching (OT-CFM) [15], Schrödinger-bridge conditional flow matching (SB-CFM) [15], as well as traditional methods scVelo [5] and Diffusion Pseudotime [2].

TrajectoryNet uses a continuous normalizing flow modeled with Neural ODE to infer the dynamic distribution of the cells. OT-CFM and SB-CFM require pre-defining conditional paths and the coupling between successive observations, and they provide two sets of couplings and conditional paths based on optimal transport and Schrödinger-bridge (marginal) methods. CFM tends to infer straight-line trajectories in Euclidean space and cannot adapt to the dynamics of cellular data, while Schrodiner bridge methods simply perform optimal transport.

In Figure 2 we visualized the inferred trajectories of our method, BEMIOflow, compared to TrajectoryNet, OT-CFM, SB-CFM.

BEMIOflow infers consistent single-cell level trajectories that provide fundamentally different analysis than widely-used methods like scVelo, which produces local arrows on the plot, but not integrated trajectory paths for each cell, and Diffusion Pseudotime, which organizes all cells into a single trajectory. Furthermore, scVelo suffers from contradictory trajectory segments, while Diffusion Pseudotime fails to produce pseudotime for cells with gaps from the starting cell. Additionally, BEMIOflow outperforms TrajectoryNet and other flow-based models, adhering better to the cellular manifold qualitatively.

In Figure 2b, we compared the root mean square error (RMSE) between ground truth and predicted trajectories in the PHATE space and gene space. Our method consistently outperformed other methods.

In addition, we computed Earth Mover’s Distance (EMD) between held-out populations and predicted populations using the trajectory inference methods. Our method consistently outperforms competing methods, demonstrating the accuracy of population interpolation.

In Figure 2c, we further compare the accuracy of gene regulatory network inference using graph edit distance over synthetic graphs in three different sizes, comparing Granger Causality (GC) (ours) with Optimal Causation Entropy (OCE) [16], PC algorithm (PC) [17, 18], multivariate Transfer Entropy (mTE) [18], multivariate Mutual Information (mMI) [19], Neural Relational Inference (NRI) [20], Diffusion Convolutional Recurrent Neural Network (DCRNN) [21], Discrete Graph Structure Learning (GTS) [22], and Dynamic Neural Relational Inference (NIR) [23].

Our method, Granger Causality (GC), largely outperforms static-expression-based methods such as OCE, PC, mTE, and mMI by directly leveraging the inferred temporal gene trajectories rather than relying on static or instantaneous gene-gene dependencies. These traditional methods fail to account for the inherent time-lagged nature of transcriptional regulation, and consequently infer noisier and less accurate networks.

Moreover, among dynamic-trajectory-based methods, including NRI, DCRNN, GTS, and NIR, GC still yields the lowest graph edit distance across all regimes. This is because GC explicitly models lagged predictive influence between genes using interpretable autoregressive models, whereas the deep learning methods either rely on latent embeddings (e.g., NRI) or require known or soft adjacency priors (e.g., DCRNN, GTS). GC, in contrast, infers directional causal edges directly from the decoded gene dynamics obtained via BEMIOflow, with no need for prior graph assumptions or sampling from latent representations. This interpretable, statistically grounded approach proves more effective for identifying causal regulatory relationships, especially in low-data and noisy biological regimes, as reflected in the synthetic benchmarks.

Uniquely, Cflows allows integration of evidence-based pruning of gene regulatory relationships. This modality is highly customisable and can use experimentally derived regulatory or gene-regulatory canvases from established knowledge bases. Using tools like Cytoscape, it is possible to customise multiple layers of such gene-regulatory or gene-interaction datasets. Next, by embedding results from Granger causality analysis, users can map and explore regulatory relationships or interaction hubs, curated with known interactions reported in the literature, as well as explore unexplored gene-relationships modules in the context of time-lapsed data. In this study, we use the TRRUSTv2 db, a gene-regulatory relationship database (https://www.grnpedia.org/trrust/), to identify temporal gene regulatory interactions of TNBC through the process of tumor formation.

#### In vitro tumorsphere data generation for exploration of trajectories and tumor-initiating cells

Tumors comprise heterogeneous cancer cell populations, of which cancer stem cells (CSCs) are uniquely endowed with the ability to initiate a tumor from a single cell (Figure 3a-b). CSCs are known drivers of cancer metastasis and are inherently resistant to many standard therapies. Accordingly, CSCs are clinically associated with aggressive tumor types and poor patient outcomes. To date, the identity of CSCs, and the dynamic gene regulatory networks that enable them to initiate and drive tumor progression remain elusive. Solving these challenges will greatly improve our understanding of tumorigenesis, and potentially lead to new therapeutic strategies to eradicate CSCs and curb tumor progression.

**Figure 3:**
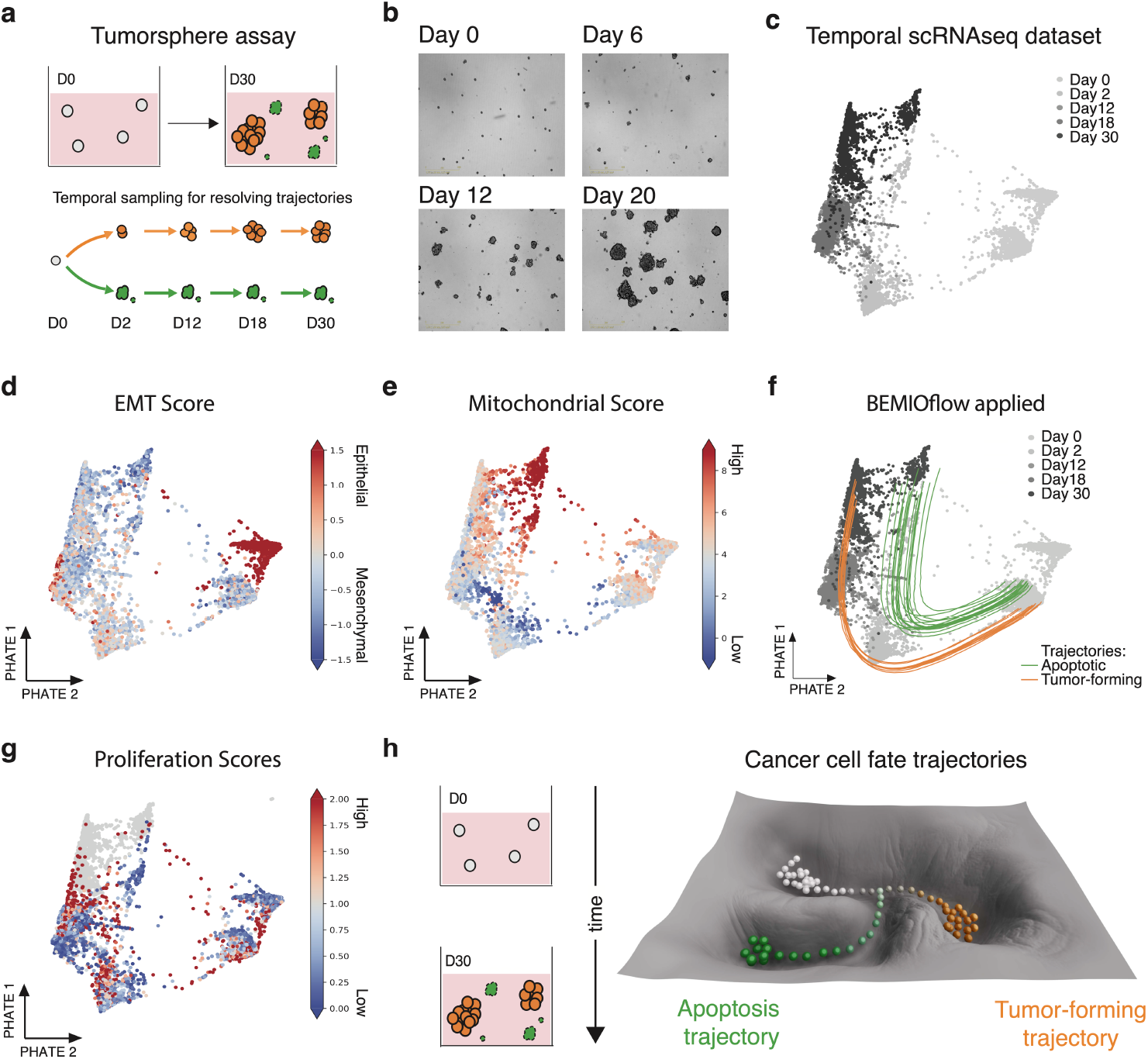
BEMIOflow applied to temporal tumorsphere scRNA-seq dataset reveals tumor-forming and apoptotic trajectories. **a,** Illustration of the tumorsphere protocol and single-cell RNAseq experiment. A heterogeneous population of triple-negative breast cancer cells (containing CSCs and non-CSCs) is seeded in a single cell suspension at day 0. By day 30, ten percent of single cells seeded produce three-dimensional heterogeneous tumorspheres. **b,** Representative images of tumourspheres across Days 0, 6, 12, 20. **c**, PHATE [12] embedding of time-lapsed scRNA-seq data generated by the tumorsphere assay described in panel a. Samples are colored by time point of data acquisition (grey scale). PHATE projections of **d**, Epithelial-to-Mesenchymal Transition (EMT) score [24] **e,** Mitochondrial activity score. **f**, Trajectories are inferred by BEMIOflow. **g,** BEMIOflow inferred proliferation scores, highlighting functional heterogeneity and progression across the experimental timeline **h,** Schematic illustrating dynamic transitions of highly tumorigenic cancer stem cells (CSCs) into two distinct cell fates through a tumor-forming or apoptotic trajectory.

While single-cell technologies have provided valuable snapshots of gene expression at different stages of tumor development, those snapshots are static and fail to capture the continuously evolving transcriptional networks over time. Moreover, computational approaches to seamlessly connect evolving cancer cell states into pathogenic trajectories and define their underlying causative gene regulatory networks remain underdeveloped.

To resolve the trajectories and dynamic molecular programs governing CSC biology, we modeled tumor formation using the 3D tumorsphere-forming assay – a surrogate *in vitro* assay for *in vivo* tumor initiation and growth [25]. In this assay, cancer cells are suspended in an anchorage-independent environment, in which only CSCs are capable of surviving and initiating a tumorsphere (Figure 3a-b). The triple-negative breast cancer cell line HCC38 was suspended as single cells in 3D, and tumorspheres emerged over a 30 day time period. Accordingly, we generated a comprehensive dataset monitoring tumorsphere-forming dynamics by performing single-cell RNA sequencing at five time points: days 0, which were the 2D cultured cell line at the point of seeding the assay, and then at days 2, 12, 18, and 30 of growth in the tumorsphere assay (Figure 3a-c). This dataset encompasses 16,983 genes and 17,983 cells (Figure 3c).

To visualize the static snapshots of tumorsphere formation we projected the 5-time points of single-cell RNA sequencing data on a PHATE space (Figure 3c). Initial analysis revealed a curved data manifold where cells organized chronologically along a clear direction of progression (Figure 3c). We used an EMT (epithelial-to-mesenchymal transition) metric score to perform an initial characterization of cell states (Figure 3d). CSCs are known to reside in a hybrid epithelial/mesenchymal state compared to their epithelial counterparts, thus the EMT scores [24] allowed us to visualize where those different cell states reside on the PHATE manifold. This embedding shows a clear distinction between non-CSCs (red dots, high EMT scores - epithelial state) and CSCs (light blue dots; lower EMT score - hybrid E/M status) on the PHATE manifold at day 0. As the tumorspheres grow, both mesenchymal and epithelial cell states emerge from day 2 to day 30.

Next we visualized mitochondrial gene expression (Figure 3e) on the PHATE manifold. Cells with high expression of mitochondrial genes (red) indicate cells likely undergoing apoptosis. At day 0, cells with high and low mitochondrial scores are evident in both the epithelial non-CSC fraction and the hybrid E/M CSC fraction. At day 2, there is an enrichment of low mitochondrial scoring cells, highlighting the population of viable cells initiating the tumorsphere. As the tumorspheres grow over the 30 time period, a population of viable cells continues to grow (blue dots), and a population of high mitochondrial scoring cells (red dots) emerges, indicating cells that are undergoing apoptosis.

#### *Cflows* refines a marker profile to isolate CSCs from a heterogeneous cell population

We applied BEMIOflow to the 5-timepoint tumorsphere formation dataset described in the previous section. BEMIOflow uncovered two distinct groups of trajectories: one terminating in dying cells with high mitochondrial scores (apoptotic trajectories), and one terminating in proliferating cells with high EMT scores (tumor-forming trajectories) (Figure 3e-f).

To obtain these trajectories, we leveraged the proliferation network component of BEMIOflow, which initially identified two distinct Day 0 subpopulations: one exhibiting high proliferation rate and corresponding to the hybrid E/M phenotype, and another with low proliferation rate (Figure 3g). We restricted trajectory inference to the high proliferation rate subpopulation to focus on cells most likely to contribute to tumorsphere formation.

Using these filtered Day 0 cells as inputs to BEMIOflow resulted in the two distinct trajectories described above. To identify the origin of the tumor-forming trajectories, we isolated BEMIOflow trajectories terminating in the tumor-forming region and identified their Day 0 starting points. We then performed kernel density estimation (KDE) on the PHATE embedding of these starting points, revealing a spatially distinct Day 0 subregion enriched for cells initiating the tumor-forming trajectory (Figure 4a), and clearly separated from the origins of apoptotic trajectories.

**Figure 4:**
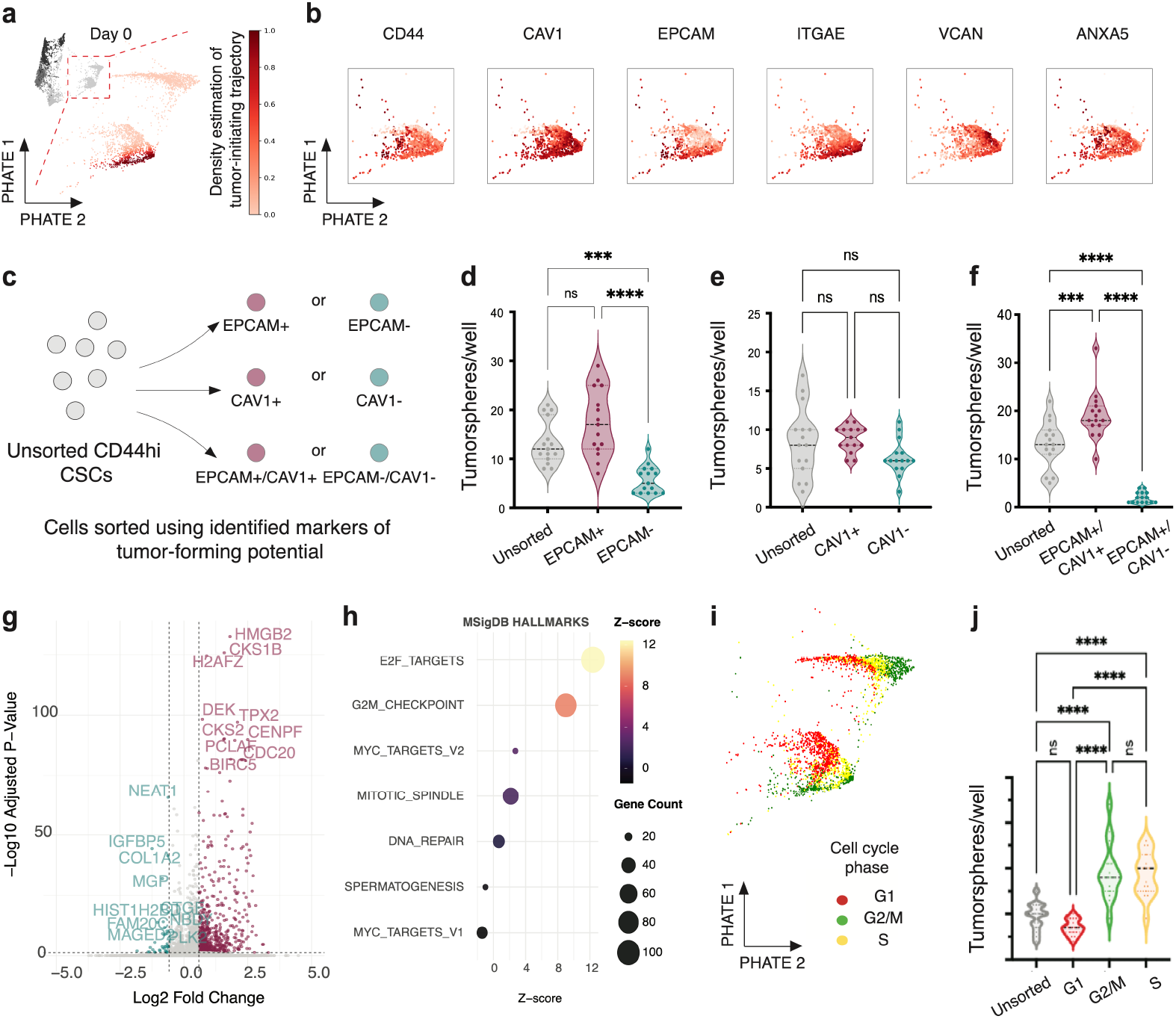
Cflows improves identification of tumor cell-of-origin. **a**, Visualization of density estimation for tumor forming trajectory origin at day 0 (inset). **b**, Visualization in CD44hi CSC fraction at day 0 of key differentially expressed cell surface markers **c**, Scheme for flow cytometry-based isolation of EPCAM+/-, CAV1+/-and double EPCAM+/CAV1+ and EPCAM-/CAV1-cells. **d-f**, Tumor-initiating potential of isolated single EPCAM+/-or CAV1+/-cells, or double EPCAM/CAV sorted cell populations (n=3, one-way ANOVA, mean +/-SEM, Tukey’s multiple comparison test) **g**, Volcano Plot showing differentially expressed genes between CD44hi Cav1hi population and CD44hi Cav1lo population with top 10 up and downregulated genes annotated **h**, Results from GSEA Hallmark pathway analysis of significantly differentially expressed genes (DEG) (adj. p-value <0.05) between CD44hi Cav1hi population and CD44hi Cav1lo population. Dot plots representing hallmark pathways with annotated z-scores and gene count (number of DEGs in pathway). **i**, PHATE plot visualizing inferred cell cycle state of cells at Day 0. **j**, Flow cytometry-based sorting of CSCs (HCC38 cells) expressing the FUCCI cell cycle sensor system to isolate cells at different stages of the cell cycle. Each cell cycle isolate is measured for *in vitro* tumorsphere-initiating potential(n=3, one-way ANOVA, mean +/-SEM, Tukey’s multiple comparison test)

Next, we identified cell surface markers highly expressed in the cells that overlap with the density of tumor-forming trajectory initiation at day 0 (Figure 4b). Notably, caveolin 1 (CAV1), and epithelial cell adhesion molecule (EPCAM) were significantly enriched and restricted to that area of the PHATE plot (Figure 4b). ITGAE (CD103), a membrane protein commonly associated with T-cell mediated tumour response and associated with CSC exosomes, was also enriched in this region of the plot [26, 27]. For comparative purposes, ANXA5 and VCAN are not specifically enriched in that area. Those data suggest that the combination of CD44, EPCAM and CAV1 or ITGAE could improve CSC isolation from a heterogeneous cell population.

To experimentally test that, we isolated EPCAM^+^ single-positive cells, CAV1^+^ single-positive cells, and EPCAM^+^CAV1^+^ double-positive cells from sorted CD44^*hi*^cells and performed the tumorsphere assay (Figure 4c). Neither single EPCAM^+^ or CAV1^+^ cells were enriched for tumorsphere-forming ability compared to unsorted CD44^*hi*^cells (Figure 4d-e). In contrast, double-positive EPCAM^+^CAV1^+^ cells had significantly increased tumorsphere-forming ability compared to CD44^*hi*^cells (1.7 fold; p*<*0.05) (Figure 4f). Thus, *Cflows* refined a cell surface marker profile to improve CSC identification and isolation.

To identify the molecular pathways driving enhanced tumor-initiating potential in the CD44^*hi*^EPCAM^+^ CAV1^+^ populations, we partitioned the CD44^*hi*^cells within the Day 0 PHATE plots based on CAV1^+^ density estimation. This allowed us to identify the differentially expressed genes between CD44^*hi*^EPCAM^+^ CAV1^+^ and CD44^*hi*^EPCAM^*lo*^CAV1^*lo*^population (Figure 4g). We performed GSEA analysis on the significant DEGs identified in these two populations (https://www.gsea-msigdb.org/gsea/msigdb/index.jsp). Hallmark signatures from MSigDB (Molecular Signatures Database) revealed CD44^*hi*^EPCAM^+^CAV1^+^ populations to be enriched for cell-cycle, including G2M check-points, DNA-repair, mitotic spindle formation among others (Figure 4h). This prompted us to perform a cell cycle analysis to computationally predict the cell cycle phase at Day 0 [28]. This analysis shows that CD44^*hi*^EPCAM^+^CAV1^+^ tumorsphere-initiating cells reside in the G2/M phase, followed by S phase (Figure 4l). To experimentally validate that finding, we expressed the FUCCI cell cycle sensor system in HCC38 cells, isolated CSCs from each cell cycle phase by flow cytometry [29], then tested each population (Unsorted, G1, G2/M or S phase) for tumorsphere-forming potential. As predicted by BEMIOflow, CSCs that reside in the G2/M and S phase are significantly enriched for tumorsphere initiation potential compared to unsorted CSCs or CSCs in the G1 phase of the cell cycle (Figure 4i).

In summary, we found a new surface marker combination that improves CSC identification and isolation, and revealed new biolgoy related to cell cycle phase that enhances tumor-initiating potential.

#### *Cflows* reconstructs cell fate trajectories and causal gene regulatory networks driving tumor-formation

As BEMIOflow learns continuous gene expression dynamics, we sought to define the causal gene regulatory networks driving the pathogenic tumorsphere-forming trajectories forward (Figure 5).

**Figure 5:**
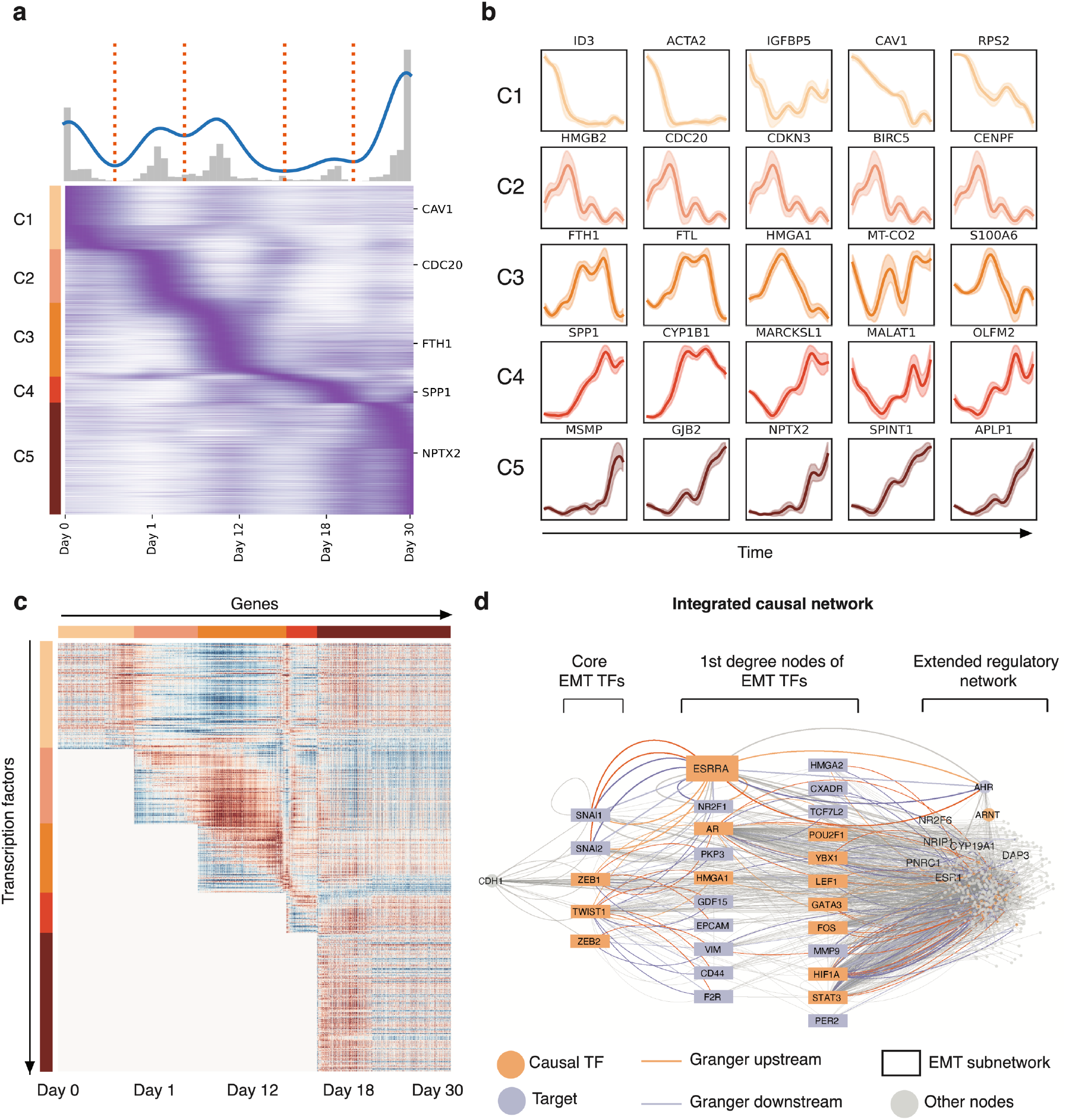
Defining the dynamic molecular program underlying tumor formation. **a**, Visualization of inferred continuous gene expression dynamics over time. Genes have been clustered into 5 groups based on their gene expression dynamics (see Methods). Key representative genes from each cluster (CAV1, CDC20, FTH1, SAT1, NPTX2) are indicated on heatmap. **b**, Visualizing inferred continuous gene expression dynamics of representative genes within each cluster. **c**, Heatmap of signed Granger values between 5,273 genes and 461 transcription factors, organized by gene clusters from Figure 5**a** for tumor-forming trajectories. Red denotes a strong enhancing relationship between a transcription factor and a target gene, while blue denotes a strong repressive relationship. **d**, Visualization of Granger-inferred transcription factor (orange nodes)–target gene (purple node) pairs driving tumor formation embedded over a regulatory network derived from TRRUST v2 database (Transcriptional Regulatory Relationships Unraveled by Sentence-based Text mining [30]). ESRRA, a causal transcription factor was associated with a subnetwork of genes including key EMT transcription factors SNAI1/2, ZEB1, CDH1 among other transcription factors including AHR and ARNT.

BEMIOFlow first decodes the inferred tumorsphere-forming trajectories back to the gene space. In Figure 5a, we show a heatmap of the mean gene expression dynamics for each gene over time, computed across all cells along the tumorsphere-forming trajectories. Genes are ordered by the timing of their peak expression, from early to late, and subsequently partitioned into five clusters based on those peak times.

We further illustrate the dynamics of representative single genes from each cluster, where the solid line represents the mean and the shaded region indicates the mean ±1 standard deviation (Figure 5b). Examples of divergent peak gene dynamics from each cluster (CAV1, CDC20, FTH1, SAT1 and NPTX2) are highlighted on the heatmap.

To uncover the temporal gene regulatory network governing the tumor-forming trajectories, we selected the top 5,273 genes with the highest and most variable expression (above the fiftieth percentile in dispersion and expression scores), as well as the top 461 variably expressed transcription factors meeting the same criteria, and performed a Granger causality analysis for every transcription factor-gene combination in the tumor-forming trajectory (Figure 5c). To ensure biological plausibility, we restricted the analysis to pairs in which the causal transcription factor belonged to a cluster that peaked earlier than the cluster of its target gene, based on the assumption that genes can only be regulated by transcription factors that peak earlier.

Given the known role of the epithelial-to-mesenchymal transition (EMT) in breast cancer biology [31, 32], we sought to reveal the temporal regulation of causal EMT drivers throughout tumorsphere formation for the first time. Using EMTdb as a reference, we selected the EMT associated genes as seed genes. We, next, filtered Granger-predicted interactions to include those where seeds were either causal regulators or downstream targets within two degrees of connectivity. These filtered gene pairs were compared to TRRUSTv2 regulatory interactions, revealing 195 unique matches. We embedded these 195 Granger causality pairs onto the TRRUSTv2 network to construct an integrated regulatory network (Figure 5d), which revealed a well-connected EMT core module comprising ZEB1/2, TWIST, and SNAI1/2. The GC analysis called ZEB1/2 and TWIST causal regulators of the tumor-forming trajectory, and SNAI1/2 as downstream targets. To identify regulatory modules around this EMT core, we projected to their first neighbors within this network and discovered that ESRRA is defined as a causal driver upstream of SNAI/1/2, and regulator of a subnetwork comprising AHR, ARNT and CDH1.

ESRRA is an orphan nuclear receptor and has recently been associated with adverse outcomes in triple-negative breast cancer [33]. It is an important mediator of cellular metabolism, and the EMT via SNAI1 and SNAI2 [34–38]. Notably, the *Cflows* ESRRA sub-network contained four causal transcription factors, including, ARNT, SNAI2, ZEB1, ZEB2 (Figure 5d). We therefore reasoned that ESRRA could be a critical driver of tumorsphere formation (Figure 5d).

#### An ESRRA-dependent gene regulatory network is essential for tumor initiation and growth

To determine if an ESRRA dependent subnetwork plays a causal role in tumorsphere formation, we first analyzed the temporal expression of ESRRA protein along with a CSC marker ZEB1, and a non-CSC epithelial marker E-cadherin, at four timepoints of tumorsphere formation (Days, 7, 14, 21 and 28; Figure 6a). This confirmed a critical temporal pattern of ESRRA expression linked to tumorsphere development. Initially, low ESRRA levels co-existed with ZEB1 at day 7, indicating co-expression in the CSC population. During days 7-14, ESRRA expression significantly increases. It then undergoes a rapid decline from day 14 onward. Notably, the down-regulation ZEB1 protein occurs after the peak in ESRRA expression at day 14, at which point the epithelial marker E-cadherin (CDH1 gene) begins to increase, indicating the emergence of the epithelial cell state (Figure 6a). Those protein expression patterns confirmed the temporal single cell gene expression patterns inferred by *Cflows* (Figure 6a-b).

**Figure 6:**
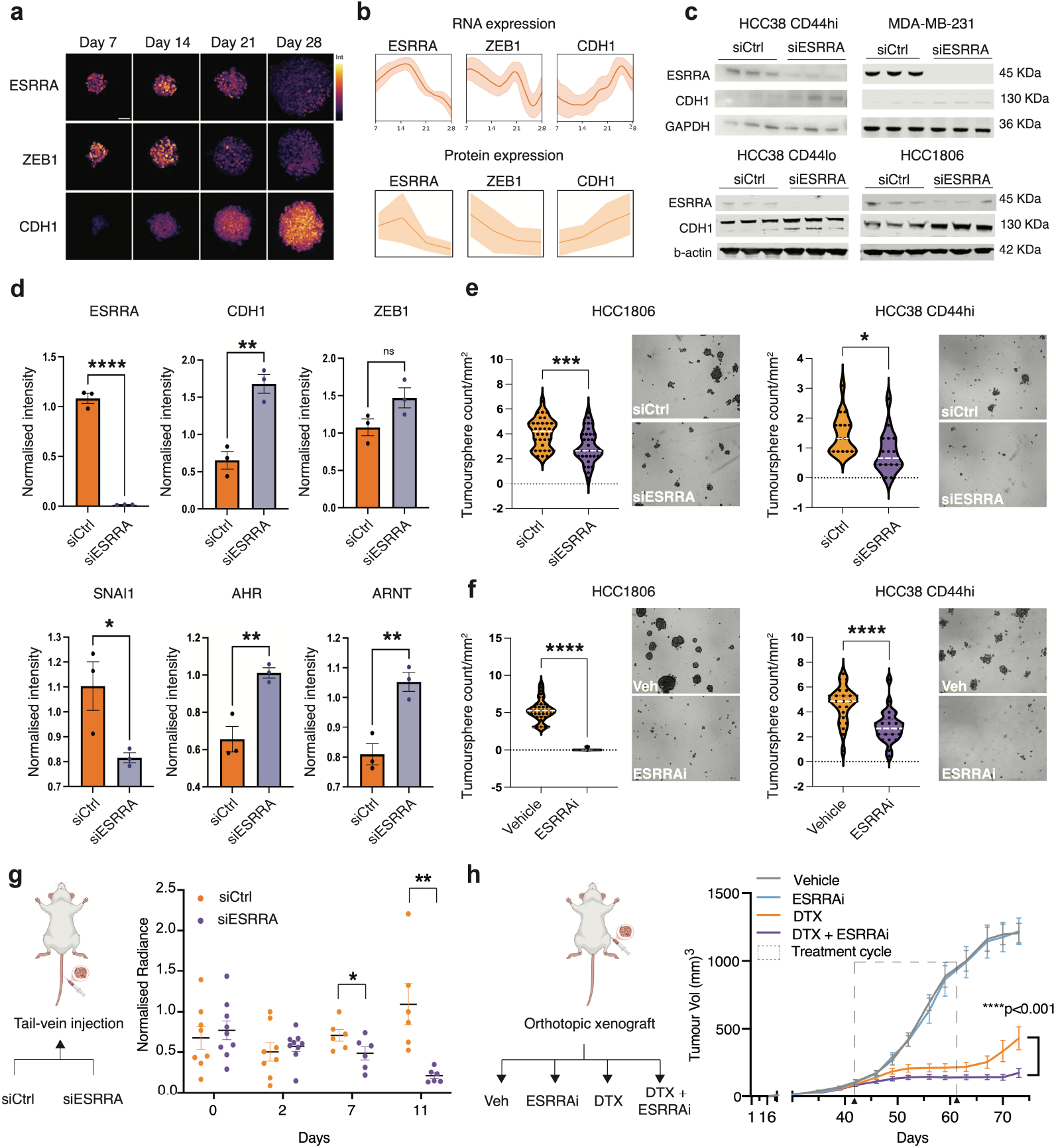
Validation of ESRRA-dependent gene regulatory network driving tumor formation. **a**, Visualization of ESRRA, ZEB1 and CDH1 by immunofluorescence staining at 4 timepoints in 3D tumorspheres. **b,** Visualization of *Cflows* interpolated gene trends as well as ground truth protein trends for ESRRA, ZEB1 and CDH1. **c,** Effect of ESRRA knockdown (siRNA) on CDH1 expression in HCC38 CD44hi, MDA-MB-231 (CSC) and HCC38 CD44lo and HCC1806 (non-CSC) cells by western blot (n=3, mean +/-SEM, students t-test, two-sided, p<0.05) **d,** Effect of ESRRA knock-down on key genes within ESRRA subnetwork by western blot (n=3, mean +/-SEM, students t-test, two-sided, p<0.05). **e,** Tumorsphere assays assessing tumor-initiating potential in control and ESRRA knock-down TNBC cell lines (HCC1806 and HCC38 CD44hi) with images of wells showing representative tumorspheres in each condition (n=3 biological replicates, n= 16 (HCC38) and n=34 (HCC1806) technical replicates per condition Student’s t-test, *p<0.05, ***p<0.001). **f,** Tumorsphere assays assessing tumor-initiating potential in vehicle and ESRRA inhibitor (ESRRAi) treated TNBC cell lines (HCC1806 and HCC38 CD44hi) with images of wells showing representative tumorspheres in each condition (n=3 biological replicates, n= 24 (HCC38) and n=36 (HCC1806) technical replicates per condition Student’s t-test, ****p<0.0001). **g,** (Left) Schematic illustrating validation strategy to test if perturbation of ESRRA affects metastatic colonisation in lungs. siControl and siESRRA MDA-MB-231 luc cells were injected via tail vein. Lung colonisation and tumor initiation was monitored using in-vivo bioluminiscence imaging. (Right) Temporal changes in normalized fold change radiance at Day 0, 2, 7 and 11 in mice injected with siControl and siESRRA MDA-MB-231-luciferase cells (Student’s t-test *p<0.05, **p<0.001) **h,** (Left) Schematic of experiment showing orthotopic MDA-MB-231 luc xenografts treated with vehicle, ESRRA inhibitor (ESRRAi), docetaxel (DTX) and combination of ESRRAi with DTX. Primary tumor growth dynamics in the MDA-MB-231 xenograft model with 1 day priming with anti-ESRRA drug (C14) before receiving chemotherapy (DTX), followed by 3 days of treatment with anti-ESRRA drug. Treatment groups include Control (Vehicle control), C14 (25mg/kg), DTX (20mg/kg) and a combination arm of DTX + C14 (20mg/kg DTX + 25mg/kg C14). This dosing schedule shows an improvement of the DTX+C14 arm (blue) in comparison with DTX (purple) with sustained and significant control of tumor volume in DTX+C14 arm at Cycle I (n= 1, 12 animals/group, mean SEM, 2-way ANOVA, Tukey’s multiple comparison test, p<0.001 day 70 onwards).

To test the causal relationship of ESRRA in controlling the CSC state, we tested the impact of ESRRA down-regulation (siESRRA) compared to control conditions (siCtrl) in multiple CSC models (HCC38 CD44^*hi*^, HCC1806 and MDA-MB-231) and a non-CSC model (HCC38 CD44^*lo*^). siESRRA led to an increase in E-cadherin (CDH1 gene) expression across all four cell lines (Figure 6c and Figure S2), which is consistent with previous work showing that ESRRA can regulate the EMT in [39–41]. We show that E-cadherin up-regulation likely results from down-regulation of the EMT transcription factor SNAI1/2 rather than ZEB1, consistent with interactions revealed by the ESRRA subnetwork and causal predictions from *Cflows* (Figure 6d). *Cflows* also predicted ARNT as a causal node. Through the network, we can visualise that ESRRA may indirectly regulate ARNT via AHR during tumorsphere formation. Indeed, ESRRA knockdown also caused significant increases in AHR and ARNT (Figure 6d-e), most widely known for their role in regulating xenobiotic metabolism as well as immune signaling and cell differentiation. Thus, the Cflows pipeline leverages knowledge bases to identify context-specific, tightly linked causality networks relevant to the biological observation. In this instance, focusing on regulatory relationships of EMT transcription factors revealed underexplored roles of ESRRA, AHR, and ARNT in tumor formation that emerge as part of a subnetwork rather than as isolated entities. Thus, while EMT drivers such as ZEB1 and SNAI1/2 are well-established mediators of tumor progression and therapy resistance [42, 43], they remain challenging to target directly [44]. Here, *Cflows* elegantly addresses this problem by enabling the identification of alternative, targetable nodes within broader regulatory subnetworks.

Next, we assessed the functional impact of ESRRA on *in vitro* tumorsphere formation using siRNA or a drug (2-Aminophenyl)(1-(3-isopropylphenyl)-1H-1,2,3-triazol-4-yl)methanone, referred to as C14 from here on) that antagonizes ESRRA function [45]. (Figure 6 f). Tumorsphere formation and size were significantly decreased in response to siESRRA as well as C14 (Figure 6f). These results confirm that ESRRA plays a critical role in tumorsphere initiation by CSCs.

To show that ESRRA is required for metastasis, ESSRA was knocked down in MDA-MB-231-*luc* cells using siRNA. Control (siCtrl) or ESRRA knockdwon cells (siESRRA) were seeded in the lungs of NOD/SCID mice via tail vein injection (Figure 6f-h). Metastatic growth in the lung was measured using bioluminescence imaging (IVIS). Metastatic growth was significantly reduced from day 7 onwards in siESRRA lungs compared to siControl lungs (Figure 6g). These data show that ESRRA is essential for CSCs to generate metastases.

Another biological attribute of CSCs is their resistance to conventional chemotherapies. Therefore, we reasoned that inhibiting ESRRA may sensitize tumors to chemotherapy. To test this, MDA-MB-231 CSCs were implanted orthotopically into the mammary fat pad of NSG mice. Once tumors reached 200 mm^3^, animals were randomized to the control arm (vehicle), C14 alone (ESRRA antagonist), docetaxel chemotherapy alone (DTX), or a combination of DTX and C14 (Figure 6h). Indeed, the DTX with C14 combination significantly reduced tumor volume compared to DTX alone (p*<*0.0001) (Figure 6h). Together, these data underscore the pivotal role of ESRRA and its gene regulatory subnetwork in tumor formation and suggest that inhibiting ESRRA is a promising strategy to decrease tumor initiation and metastasis, while boosting sensitivity to chemotherapy.

## Discussion

Dynamic cell state evolution is a hallmark of many diseases. While learning dynamics has often been a holy grail for this type of data, previous efforts have significant shortcomings, in that they cannot learn network dynamics in high-dimensional data over extended time periods. To address this knowledge-gap, we developed *Cflows*, a pipeline that learns continuous cellular dynamics and builds rich gene-regulatory networks from static snapshots of time-lapse single-cell data. Importantly, because *Cflows* only requires time-lapsed data with a minimal time points, it can be broadly applied to any biological system for which temporal data is available. Indeed, we show that *Cflows* vastly outperforms existing state-of-the-art trajectory inference methods.

*Cflows* features BEMIOflow, an enhanced version of the trajectory inference model MIOflow [4], that generates biologically-plausible, continuous cell-state trajectories. BEMIOflow includes regularizations that approximate population-level behavior, and includes growth and death rates to better capture features inherent to biological systems. The continuous nature of these trajectories enables the use of time-based causality analysis. *Cflows* uses Granger Causality to identify causal transcriptional networks from BEMIOflow-inferred gene dynamics, which are refined using existing knowledge databases to increase trust. After inferring these networks, we validated them with targeted *in vivo* experiments.

In cancer, cell state transitions are an essential feature of tumor growth and cancer progression [46]. Here we applied *Cflows* to model the dynamic cell state transitions and gene regulatory networks that drive tumor growth originating from CSCs - a subpopulation of aggressive cancer cells with the unique ability to initiate a tumor from a single cell. While that biological property of CSCs has long been established, the ability to model the temporal gene regulatory networks that enable a single CSC to enter the cell cycle and generate a robustly growing tumor are poorly understood. Here, we show for the first time that *Cflows* revealed the rich and complex causal networks that enable those decisions to be made.

Density approximation of the origin of *Cflows* cellular trajectories enabled the identification of a cell surface marker profile to isolate CSCs from a heterogeneous population of cancer cells. We experimentally validated that flow cytometry isolation of CSCs with a CD44^*hi*^EPCAM^+^CAV1^+^ profile, and, residence in the G2/M-or S-phase of the cell cycle, significantly improved identification of CSCs with enhanced tumor-initiating ability. Thus, *Cflows* can be used to identify key cell states in heterogeneous cell populations.

We also computed continuous time-varying gene trends for every measured gene, in every cell. We then grouped the cellular trajectories into two types: tumor-forming and apoptotic. We further studied the tumor-forming trajectories by identifying 5-clusters of gene based on peak-time trends. We showed these gene trends revealing a gene expression program that consists of a cascade of genes that go up and down in sequence. These timed gene trends then allowed us to compute causal gene regulatory networks.

We performed extensive experimental validation of the *Cflows* predicted causal gene regulatory networks that enable CSCs to develop into a robustly growing tumorsphere. We identified an ESRRA subnetwork (including SNAI1, AHR and ARNT) as a critical driver of CSC biology, and experimentally validated that ESRRA inhibition drives CSC into a nonCSC state *in vitro* and *in vivo*. Moreover, we show that the inherent chemotherapy-resistant phenotype of CSCs is significantly reduced by inhibiting ESRRA in combination with chemotherapy. Together, these data suggest that ESRRA-targeted therapies could be an effective clinical strategy to curb CSC-dependent tumor growth, metastasis, and resistance to chemotherapy [46].

The *Cflows* pipeline is a breakthrough method that can unravel complex temporal transcriptional networks governing cell state dynamics. This pipeline can be used to study any type of dynamic transition captured via single cell technology over multiple timepoints, and may include stem cell differentiation, response to therapeutic interventions, or infections. As longitudinal data becomes increasingly available and critical in the study of dynamic systems, *Cflows* will be applied to a broader set of datasets and types.

## Author Contributions and Acknowledgements

Smita Krishnaswamy and Christine Chaffer conceived the project. Smita Krishnaswamy, XIngzhi Sun and Alex Tong developed the Cflows algorithm. Xingzhi Sun, Shabarni Gupta, Alexander Tong, Manik Kuchroo and Dhananjay Bhaskar jointly led this work under the supervision of Smita Krishnaswamy and Christine Chaffer, who were collectively responsible for writing the manuscript. Smita Krishnaswamy, Xingzhi Sun, Dhananjay Bhaskar, Alexander Tong, Manik Kuchroo, Aarthi Venkat, and Brandon Zhu were responsible for algorithm development, running computational comparisons, and applying *Cflows* to biological datasets under the supervision of Smita Krishnaswamy. Shabarni Gupta was responsible for network building and validation, target manipulation for in vitro and in vivo experiments. Beatriz Perez San Juan was responsible for single-cell experiments, FACS analysis, and cell sorting, tumorsphere assays. Laura Rangel was responsible for tumorsphere assays and fluorescent staining, John Lock performed the analysis of immunofluorescence experiments, Vanina Rodriguez and Shabarni Gupta were responsible for monitoring and treating the animals for all *in vivo* experiments presented, all under the supervision of Christine Chaffer. Chen Liu contributed to figure preparation and assisted in writing the manuscript. Andrew Cox was also instrumental in the writing of this manuscript.

## Ethics statement

Animal experiments were conducted with approval from the Garvan Institute of Medical Research Animal Ethics Committee and in adherence to the Australian Code of Practice for the Care and Use of Animals for Scientific Purposes (Project code ARA_21_24).

## Inclusion statement

All contributors to this manuscript meet the authorship criteria mandated by Nature Portfolio journals and have been acknowledged as authors, as their participation was integral to this study. Roles and responsibilities were agreed upon among collaborators. The study highlights a scientific question of global importance and includes findings through a network of international collaborators. Each collaborator involved local partners who are involved in this study and have been included as authors. The research faced no constraints or prohibitions in the researchers’ environment, ensuring it avoided stigmatization, incrimination, discrimination, or personal risks to participants.

## Declaration of Interests

The authors declare that they have no conflict of interest.

## Methods

### Overview of *Cflows*

*Cflows* is an end-to-end computational framework for inferring gene regulatory networks from single-cell data. It integrates flow-based generative modeling, dynamic optimal transport, and causal inference. The core of *Cflows* is BEMIOflow, a trajectory inference model that builds upon our previous method, MIOflow, by introducing the ability to model both cell proliferation and death - key features in systems such as cancer. *Cflows* first uses BEMIOflow to learn continuous gene-level trajectories from snapshot data, and then applies Granger causality analysis to infer gene interactions from these learned dynamics. We describe the components of the framework below.

### Embedding Gene Expression into the Manifold of Cell States

The first step in the Cflows pipeline is to embed the single-cell gene expression data into a space that reflects the underlying structure of cellular transitions. Unlike traditional models that operate directly on raw expression values or low-dimensional PCA coordinates, Cflows uses a more biologically meaningful latent space that captures how cells evolve over time. Specifically, we embed the data into a PHATE-based manifold that preserves both the geometry and local connectivity of the transcriptomic landscape. This space forms the foundation for learning continuous trajectories with neural ODEs in the next stage of the pipeline.

We begin by reducing the dimensionality of the data using Principal Component Analysis (PCA). Given the gene expression matrix 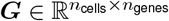, we subtract the mean expression vector to center the data:

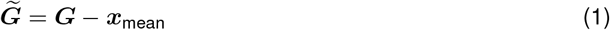

and then project it onto the top 50 principal components:

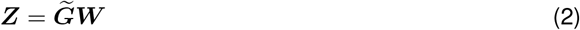

where ***W***contains the PCA loadings. This gives us an initial low-dimensional representation of each cell.

Next, we use PHATE to compute diffusion-based distances between cells in this reduced space. These “potential distances” capture how easily information can flow between different cell states, making them particularly well-suited for modeling developmental processes. To embed the data in a way that preserves these relationships, we train a neural network encoder, *h*_*ξ*_, that learns a new coordinate system in which Euclidean distances reflect PHATE distances. The encoder is trained using a distance-matching loss:

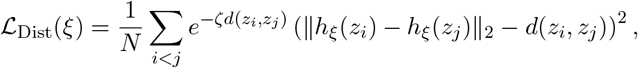

which emphasizes preserving local structure by weighting nearby cells more heavily.

The result is a latent space that captures the smooth progression of cell states along biological processes like differentiation or transformation. By learning trajectories in this space, rather than in raw gene space, Cflows can model transitions in a way that aligns with how cells actually behave—following continuous, low-energy paths through a realistic cellular landscape.

### Trajectory inference using BEMIOflow

BEMIOflow learns a neural ODE model that interpolates between cross-sectional measurements of cell populations, producing continuous trajectories that represent biologically plausible cellular dynamics. In this section, we provide the mathematical foundations and describe the training procedure for BEMIOflow. We begin by reviewing the neural ODE framework used to parameterize cell dynamics, followed by its formulation within a dynamic optimal transport problem. We then describe the loss functions used to train the model, which include trajectory smoothness, distributional matching, and manifold regularization. Finally, we introduce the main enhancement over MIOflow - the modeling of cell proliferation and death through a learned growth rate function.

### Neural Ordinary Differential Equation Model

BEMIOflow models cellular transitions as solutions to a neural ODE. A neural ODE defines a vector field *f*_*θ*_(*x, t*) parameterized by a neural network, such that the evolution of cell states *x*(*t*) over time is governed by 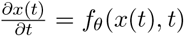. Given an initial state *x*(*t*_0_), the state at a later time *t* is obtained by integrating this vector field, 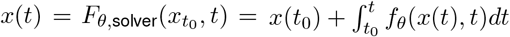.

This integral could be computed using Euler integration where at a fixed set of times *t*_0_, *t*_1_, *… t*_*T*_ we calculate 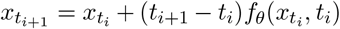 incrementally. However, this can accumulate significant error depending on the dynamics, therefore we use ODE solvers with lower error and/or adaptive step sizes.

To train the model, we compute gradients of a loss function *L* with respect to the neural network parameters *θ* using the adjoint method [7]. This involves solving an adjoint equation backward in time using the adjoint state *a*(*t*) = *∂L/∂x*(*t*), which allows us to compute *∂L/∂θ* without storing intermediate states - substantially reducing memory usage.

### Optimal Transport

Optimal transport provides a mathematical framework for moving one distribution (probability mass) to another while minimizing cost. Optimal transport considers a *ground distance* defined between points and lifts this to distances between measures. Given a metric measure space ℳ = (𝒳, *d*), the squared 2-Wasserstein distance between two distributions *µ* and *ν* on ℳ is defined as:

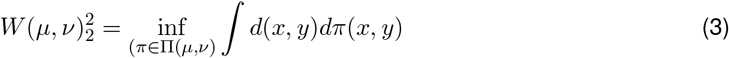

where Π(*µ, ν*) denotes the set of joint distributions (couplings) with marginals *µ* and *ν*, and both distributions are assumed to be normalized such that 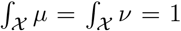. This formulation seeks the minimal total cost required to move the mass from *µ* to *ν*.

While elegant, this optimization is difficult to solve in high-dimensional continuous spaces. In the following sections, we show how *Cflows* uses dynamic optimal transport to approximate this problem.

### Dynamic Optimal Transport

Dynamic optimal transport extends the static OT formulation by introducing a time component. This makes it possible to connect OT to dynamical systems and fluid mechanics [47]. Rather than directly matching two static distributions, the dynamic formulation describes a continuous-time interpolation between them. Given a time interval [*t*_0_, *t*_1_], we seek a time-varying probability distribution *ρ*(*x, t*) and a velocity field *f* (*x, t*) such that the distribution evolves according to the continuity equation:

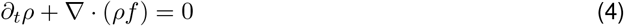

for *t*_0_ < *t* < *t*_1_ and *x* ∈ ℝ^*d*^, with the boundary conditions:

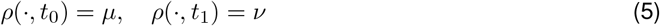

Under these constraints, the squared 2-Wasserstein distance becomes:

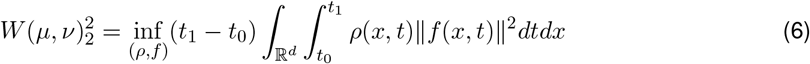

In other words, a velocity field *f* (*x, t*) with minimum *L*^2^ norm that transports mass at *µ* to mass at *ν* when integrated over the time interval is the optimal plan for an *L*^2^ Wasserstein distance. Note that the optimal paths for each point pair, (*x*_0_, *x*_1_), are geodesics and that inf_*f*_ ∥*f*_*t*_ (*x, t*) ∥^2^ = *d*(*x*_0_, *x*_1_)^2^. This problem is in general challenging to solve, particularly in the high dimensional case where a discretization of space-time (such as used in [48]) is impractical. We instead use neural ODEs to parameterize and learn the vector field *f*, which allows efficient approximation of the transport plan.

### Dynamic optimal transport with a neural ODE

Using a neural ODE, we parameterize a time-varying vector field that generates transport trajectories. Instead of enforcing hard boundary constraints *ρ*(·, *t*_0_) = *µ* and *ρ*(·, *t*_1_) = *ν*, we keep the constraint at *t*_0_ and replace the one at *t*_1_ with a relaxed penalty term, *D*(*ρ*(·, *t*_1_), *ν*), where *D* is a dissimilarity function that vanishes if and only if *ρ* = *ν*. This relaxation allows for tractable optimization using deep learning methods. The following theorem from [4] formalizes this connection:

#### Theorem 1

(Theorem 3 from [4]). *Let f* (*x, t*) *be a time-varying vector field defining trajectories dX*_*t*_ = *f* (*X*_*t*_, *t*)*dt with density ρ*_*t*_ = *ρ*(·, *t*). *Let D*(*µ, ν*) = 0 *iff µ* = *ν. Then, there exists a sufficiently large λ* > 0 *such that:*

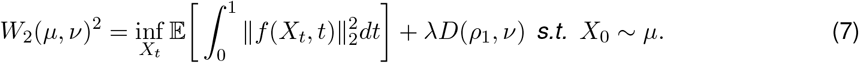

By parameterizing our field *f* with a neural network, this theorem allows us to optimize for the dynamic optimal transport flows with deep learning techniques. For a full proof see [4], we provide a proof sketch here. This result can be seen by viewing *λ* as a Lagrange multiplier over the relaxing the constraint *ρ*_1_ = *ν*. As *λ* → ∞, the relaxation enforces *ρ*_1_ ≈ *ν*. We refer the reader to [3, 4] for further details.

### Enforcing transport on a manifold

It is widely accepted that high-dimensional data often concentrate near a low-dimensional manifold, a concept generally known as the manifold hypothesis [49–51]. Manifold learning methods such as Diffusion Maps [52], PHATE [12], GAGA [13], CUTS [53], DSE [54], DYMAG [55], DiffKillR [56] and HeatGeo [57] exploit this concept and use diffusion-based affinities to recover the underlying geometry, enabling robust analysis even in the presence of noise and sparsity.

As illustrated in Supplementary Figure S1, BEMIOflow constrains transport to lie on a data manifold learned using PHATE [12], which captures the space of biologically realistic cell states. This is achieved by a PHATE-regularized encoder-decoder model [13] whose latent space resembles the PHATE space.

To ensure that the predicted trajectories remain faithful to the observations within this manifold, we we use a density regularization loss:

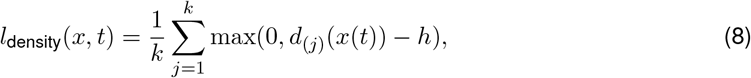

where *x*(*t*) is the point on the trajectory at time *t, d*_(*j*)_(*x*) is the distance to the *j*^*th*^ nearest neighbor in the dataset 𝒳, and *k, h >* 0 are hyperparameters. This penalty encourages trajectories to remain within a distance *h* of their *k* nearest neighbors in the observed data.

### Loss formulation for trajectory inference

We now describe the loss functions used to train the neural ODE model underlying BEMIOflow. Cells are modeled as points in a continuous state space 𝒞 ⊂ ℝ^*d*^over a time interval 𝒯 = [0, *T* − 1]. Let ***X***= {*X*_*i*_ : *i* = 0, *…, T* − 1} denote the observed cell populations at discrete timepoints. Each *X*_*i*_ contains *n* observations 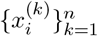 in some space 𝒳. Because single-cell measurements are destructive, each cell is observed at only one timepoint.

To improve trajectory inference in the presence of cellular heterogeneity within individual timepoints, we apply an intra-timepoint balancing procedure as a pre-processing step. Specifically, we cluster cells within each timepoint based on their PHATE coordinates, uniformly resample an equal number of cells from each subcluster to avoid overrepresentation of dominant cell states, and refine their relative positions using local pseudotime ordering. This strategy improves the continuity of inferred dynamics by encouraging the ODE to capture internal structure within each timepoint, without influencing downstream estimation of growth or death rates, which is handled independently by the proliferation network described later.

We assume that cells evolve evolve according to an ODE, *dx*(*t*) = *f* (*x*(*t*), *t*)*dt*, where *x*(*t*) ∈ 𝒞 denotes the state of a cell at time *t*, and *f* is a time-varying vector field. We parameterize *f* as a neural network *f*_*θ*_, and generate trajectories by integrating from an initial population 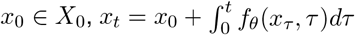. We denote the predicted cell states at time *t* as 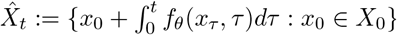. Let 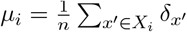 be the empirical distribution of observed cells at time *t*_*i*_, where the delta function 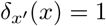 if *x* = *x*^′^ and 0 otherwise. Likewise, let 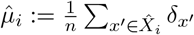 be the empirical distribution of the set of predicted points 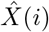.

To train *f*_*θ*_, we minimize a weighted sum of three loss components:

1. The *energy loss*, which previously appeared in the first term of Equation (7):

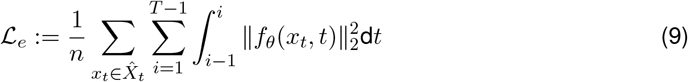
2. The *optimal transport loss*, as appeared in the second term of Equation (7), where we use 2-Wasserstein distance for the dissimilarity function *D*:

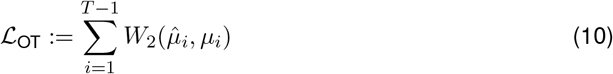
3. The *density loss* that enforces the trajectories to stay on the data manifold:

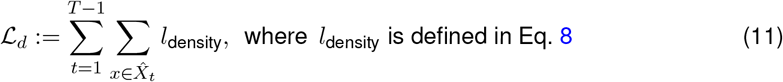

The final loss function is:

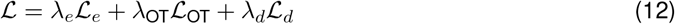

### BEMIOflow Models Growth Rate Using A Proliferation Neural Network

To model proliferation and death, BEMIOflow introduces a proliferation neural network *g*_*ϕ*_(*x, t*). For a cell at state *x* and time *t, g*_*ϕ*_(*x, t*) estimates the expected number of descendant cells at time *t* + 1. Values *m* = *g*_*ϕ*_(*x, t*) less than 1 indicate expected death, and values greater than 1 indicate proliferation. This modifies the predicted marginal distribution:

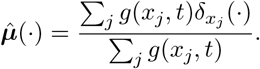

This weighted distribution is used in the marginal loss:

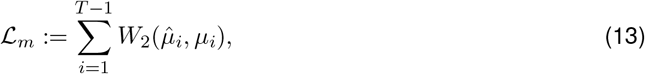

where 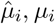 are the predicted and ground truth discrete distributions respectively with uniform weight. We initialize the proliferation network using unbalanced optimal transport [3]. Specifically, we solve:

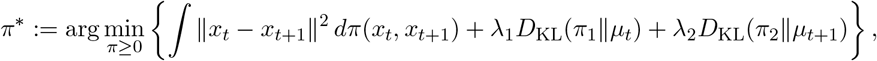

where 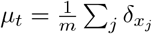 and 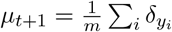. *π*_1_ and *π*_2_ are the marginals of *π*. We use a small *λ*_1_ and large *λ*_2_ to allow mass change at time *t* and encourage alignment at *t* + 1.

We estimate the initial growth function using the first marginal of *π*^∗^:

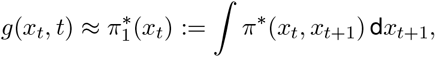

which we use to warm-start the downstream training.

### *Cflows* identifies the CSC niche using kernel density estimation

To distinguish the tumorsphere-initiating **Cancer Stem Cell (CSC)** population from the non-initiating cells within the Day 0 cohort, we developed a density-based approach. Our strategy uses the trajectory-initiating cells previously identified by BEMIOflow as a proxy for the true CSC population.

First, we built a **Kernel Density Estimation (KDE)** model using the 2D PHATE coordinates of these trajectory-starting cells ({***x***_init,*i*_}). We then evaluated this density function across all cells (*x*) in the entire Day 0 population to assign each one a “CSC likelihood” score:

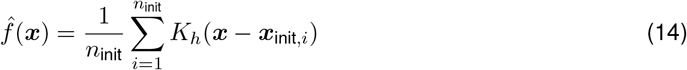

where *n*_init_ is the number of trajectory-initiating cells and *K*_*h*_ is a scaled Gaussian kernel where *K*_*h*_(*u*) = (1*/h*)*K*(*u/h*) and *K*(*u*) ∝ exp(−∥***u***∥^2^*/*2).

This KDE-based method provides a more complete picture of the CSC population by using the BEMIOflow-identified cells to create a density map, which highlights the entire neighborhood of similar, tumor-initiating cells within the PHATE embedding (Figure 4a).

### *Cflows* decodes gene dynamics and transcriptional programs from PHATE space

To recover dynamic gene expression profiles from the low-dimensional cell trajectories inferred by BEMIOflow, we developed a two-step decoding strategy. This maps the learned cellular dynamics from PHATE space back into the original gene expression space, enabling direct biological interpretation of inferred trajectories.

In the first step, we train a multi-layer perceptron, referred to as the **PCA-space decoder** *D***_*η*_**, to translate PHATE coordinates (***x*** ∈ ℝ^2^) into the 50-dimensional PCA space (***z*** ∈ ℝ^50^). The decoder is optimized by minimizing a weighted reconstruction loss that compares the decoded PCA coordinates to the original PCA embeddings of the cells:

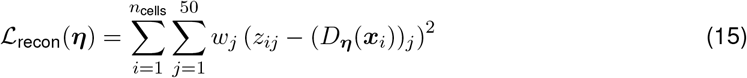

Here, 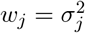 is the variance explained by the *j*^th^ principal component, emphasizing accurate recon-struction of high-variance directions in the PCA space.

Once trained, the decoder is applied to each point along a trajectory ***x***(*t*) inferred by BEMIOflow to generate a corresponding time-varying representation ***z***(*t*) in PCA space:

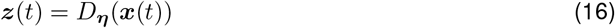

In the second step, we reconstruct gene expression profiles from these PCA coordinates using the inverse PCA transformation:

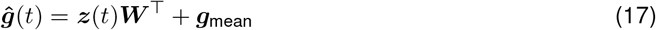

where ***W*** is the PCA loading matrix and ***g***_mean_ is the mean gene expression vector used during the initial PCA.

In practice, we discretize each continuous trajectory into 100 equally spaced timepoints to generate a gene-by-time matrix. These reconstructed expression dynamics serve as the foundation for downstream analysis, including the identification of transcriptional programs and regulatory interactions.

### Identifying Dynamic Transcriptional Programs

To obtain an overview of the transcriptional programs, we aggregate, order, and group the decoded gene dynamics.

First, to capture the aggregate behavior of each gene, we compute the mean expression dynamic across all cell trajectories. To compare the temporal patterns irrespective of differences in expression magnitude, we apply min-max scaling to normalize each gene’s mean trajectory to the range [0, 1]. For a given gene’s mean expression profile ***g***(*t*), its normalized trend 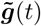 is:

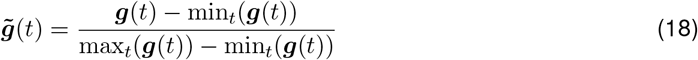

Next, we identify the **peak activation time** for each gene by finding the time point of maximum expression in its normalized trend:

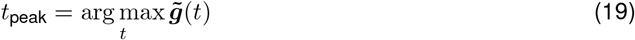

Sorting all genes by their peak time reveals a temporal cascade of gene activation (Figure 5a).

Finally, we group these ordered genes into clusters. We perform **Kernel Density Estimation (KDE)** on the distribution of the *n* peak times:

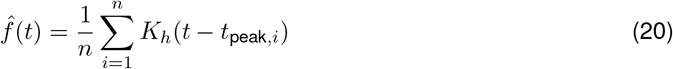

where *K*_*h*_(*u*) = (1*/h*)*K*(*u/h*) is a scaled Gaussian kernel *K*(*u*) ∝ exp(−*u*^2^*/*2). We then identify the modes of this density distribution by detecting its local minima. A point is considered a local minimum if its value is lower than all points within a surrounding window (5 points on each side). This process identified 4 local minima, partitioning the genes into 5 distinct groups (Figure 5a,b). Each cluster therefore defines a module of genes that share a similar temporal activation pattern.

### *Cflows* learns transcriptional networks from dynamic gene trends using Granger causality

*Cflows* combines continuous gene expression trends with time-lapse causality analysis as well as public databases to learn the transcriptional networks underlying cellular trajectories. *Cflows* outputs complete transcriptional trends for each cell and gene across time. We utilize this information to infer better gene-gene relationships across time using Granger causality. Granger causality goes beyond correlations in time series by testing whether a variable (or set of variables) *X* forecasts a variable *Y*. More specifically, a variable *X Granger-causes* a variable *Y* if predictions of *Y* based on its past values and on the past values of *X* are better than the predictions of *Y* based on only its own past values. To determine the likely gene regulatory structure, we use Granger causality analysis on the average *Cflows* trajectories for groups of cells. We determine that there is a directed edge from a gene *X* to gene *Y* if gene *X* Granger-causes *Y*. We score this edge based on the Granger *p*-value signed by the direction of influence of *X* on *Y*. This is determined by the sign of the coefficient on *X* in the linear regression of *Y* based on the past values of *X* and *Y*. If this coefficient is positive, we take this to mean that *X* has a likely up-regulatory effect on *Y*. Conversely, if the linear regression coefficient of *X* is negative, we take this to mean *X* has a down-regulatory effect on *Y*. By taking the log transformed Granger test statistic and adding a sign for direction of effect between a transcription factor and a target gene, we compute an interaction strength score that determines whether a transcription factor *X* may have an effect on a particular target gene *Y*. By aggregating signed Granger analysis results across large numbers of transcription factors and genes, we can identify transcriptional programs responsible for defining key trajectories.

We note that Granger causality is not the same as “true causality” as it is only evidence of *X* preceding *Y*, and there are many instances where this precedence may be incidental where changing *X* does not change *Y*. We, therefore, combine these results with known gene-gene relationships found in the Transcriptional Regulatory Relationships Unraveled by Sentence-based Text mining (TRRUST v2) database [30]. While these databases are also imperfect, the integration of trajectory inference, causality analysis and a public gene regulatory network better approximates the underlying transcriptional network. In this instance, we first identify the key driver transcription factors using a total Granger causal score (TGCS):

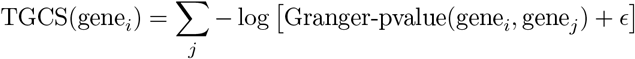

with the idea that key drivers will strongly Granger-cause many other genes. We show that TGCS outperforms other methods which assume static cellular states at inferring gene regulatory interactions from time series data (Figure 2d). Following TGCS scoring we then use the TRUSST v2 database to generate a seed network (using Cytoscape 3.9.0). We then use this seed network to extract the first-degree nodes of the top drivers of the trajectory (based on TGCS score) and their adjacent edges. We further prune and weigh the edges of this network with Granger scores to create higher fidelity networks (see methods). We do however, caution that Granger scores do not prove causality and that these should be regarded as candidates for further validation.

### Comparative methods for inferring transcriptional networks from gene expression trajectories

**Multivariate mutual information (mMI)** [19] quantifies statistical dependence between multiple genes without assuming linearity. It computes the shared information among variables *X*_1_, *X*_2_, *…, X*_*n*_ and a target gene *Y* using the multivariate mutual information:

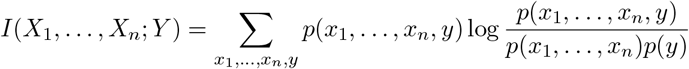

Edges are inferred between gene pairs with high mutual information, indicating strong statistical dependence.

**Multivariate transfer entropy (mTE)** [18] measures directed information transfer between time series. For genes *X* and *Y*, mTE quantifies how past values of *X* improve prediction of future values of *Y*, conditioned on other variables:

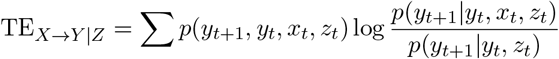

A nonzero value indicates directional influence from *X* to *Y*.

**Optimal Causation Entropy (OCE)** [16] identifies the minimal set of variables that causally influence a target by minimizing conditional entropy. For a target *Y*, the causal set *S*^∗^ satisfies:

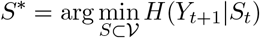

where 𝒱 is the set of all other genes. This method produces sparse causal networks.

**PC Algorithm** [17, 18] infers causal structure via conditional independence testing. It begins with a fully connected undirected graph and removes edges when conditional independence is detected (e.g., *X* ⊥ *Y* | *Z*). It then orients the remaining edges using v-structure identification and propagation rules. A v-structure occurs when two nodes *X* and *Y* both influence a common child *Z*, forming *X* → *Z* ←*Y*, and *X* and *Y* are not directly connected. Propagation rules (e.g. Meek rules) then systematically orient additional edges to avoid cycles and ensure consistency with observed dependencies, producing a partially directed acyclic graph.

**Neural Relational Inference (NRI)** [20] uses a variational autoencoder to infer latent interaction graphs from trajectories of node features. Given sequences {*x*_*i*_(*t*)}, it infers a latent adjacency matrix *A* and learns dynamics via a message-passing neural network conditioned on *A*. The model is trained by maximizing the evidence lower bound (ELBO) on trajectory likelihoods.

**Diffusion Convolutional Recurrent Neural Network (DCRNN)** [21] learns temporal dynamics on a fixed graph by modeling diffusion processes. Given a known or initialized adjacency matrix *A*, DCRNN applies diffusion convolution to node features and combines it with gated recurrent units (GRUs) to capture nonlinear temporal dependencies. The graph structure is refined through training to predict temporal dynamics.

**Discrete Graph Structure Learning (GTS)** [22] jointly learns temporal dynamics and the underlying discrete graph structure. It alternates between structure learning and time series prediction, updating a learnable adjacency matrix *A*. To sample discrete edges during training, it uses the Gumbel-Softmax trick, which provides a continuous, differentiable approximation to categorical sampling. Specifically, edge presence is modeled as:

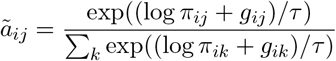

where *π*_*ij*_ is the probability of an edge from *i* to *j, g*_*ij*_ is a Gumbel noise sample, and *τ* is a temperature parameter. This enables gradient-based optimization of discrete graph structures.

### Simulation of cell differentiation with variable proliferation rates using SERGIO

We used SERGIO [10] to simulate single-cell gene expression data from ground truth transcriptional networks under three distinct scenarios: (1) a bifurcation into two terminal lineages, (2) a unidirectional linear trajectory, and (3) a cyclical process resembling periodic cell state transitions.

We first defined an underlying gene regulatory network (GRN) consisting of master regulators (MRs) and their downstream targets. In this framework, each gene *i* has a continuous expression level *x*_*i*_(*t*) governed by a chemical Langevin approximation of the chemical master equation. Specifically, the time evolution of *x*_*i*_(*t*) is described by the stochastic differential equation

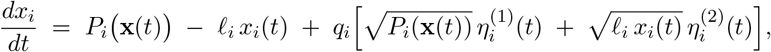

where 𝓁_*i*_ denotes the first-order decay rate, *q*_*i*_ is the noise amplitude, and 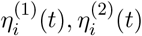 are independent Gaussian white-noise processes (mean zero, unit variance). The production rate *P*_*i*_(**x**) captures combined regulatory inputs from all parent regulators *j* ∈ *R*_*i*_ via Hill-type functions:

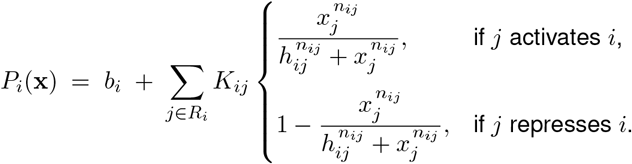

Here, *b*_*i*_ is the basal production rate (nonzero only for MRs), *K*_*ij*_ is the maximal regulatory strength, *n*_*ij*_ is the Hill coefficient, and *h*_*ij*_ is the half-saturation constant (set equal to the expected regulator steady state). MRs have no upstream regulators, so *P*_*i*_(**x**) = *b*_*i*_ for those genes. The parameters *𝓁*_*i*_, *q*_*i*_, *K*_*ij*_, *n*_*ij*_, *b*_*i*_ were selected to reflect moderate noise and biologically plausible decay dynamics; *h*_*ij*_ was computed by first estimating each gene’s steady state to initialize the system near equilibrium and minimize burn-in. SERGIO integrates these SDEs using the Euler–Maruyama scheme with a small time step Δ*t*. At each step *t* → *t* + Δ*t*, each gene *i* is updated as

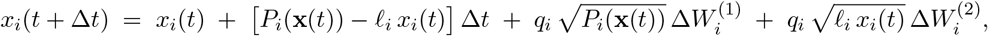

Where 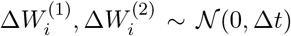. For each predefined cell type, we set the MR basal rates *b*_*i*_ to distinct values and sampled expression vectors **x**(*t*) at discrete time points to represent individual cells. After generating a continuous expression matrix 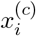 (genes × cells) for each replicate, SERGIO applied a technical-noise pipeline, including library size variation and dropout, to emulate typical scRNA-seq measurements.

In our simulations, the unidirectional and bifurcation trajectories were generated entirely using SERGIO. For the cyclic process, we used scVelo [5] to generate latent dynamics that reflect periodic transitions between cell states, since SERGIO is not designed to model cyclic attractors. However, we applied SERGIO’s noise model to the resulting expression data to simulate realistic technical effects and maintain consistency across datasets.

For the unidirectional and cyclic scenarios, cells were sampled uniformly across pseudotime, resulting in a constant density of cells along the trajectory. In contrast, the bifurcation scenario incorporated variable proliferation rates by modulating the sampling frequency of cells along the two branches. Specifically, regions with higher proliferation were sampled more densely, producing an uneven distribution of cells that mimics lineage-specific expansion. These datasets were used to evaluate *Cflows* and other computational methods.

### Other Computational Methods Details

#### Single-cell RNA sequencing and pre-processing

scRNA-seq data from 5 samples were processed with the 10X and CellRanger pipeline according to the following steps. Sample demultiplexing and read alignment to the NCBI reference GRCh38 was completed to map reads using CellRanger (v3.0.2). We prefiltered using parameters in scprep (v1.0.3, https://github.com/KrishnaswamyLab/scprep). Cells that contained at least 1,000 unique transcripts were kept for further analysis to generate a cell-by-gene matrix containing 17,983 cells and 16,983 genes. Normalization was performed using default parameters with L1 normalization, adjusting the total library size of each cell to 10,000. Any cell expressing mitochondrial genes greater than 10% of their overall transcriptome was removed. Raw data files for scRNA-seq data will be available for download through GEO under an accession number to be assigned with no restrictions on publication.

#### Transcriptional network generation

Transcriptional networks were built to visualise direct and indirect gene regulatory interactions of core transcription factors (transcription factors) identified by *Cflows* within the tumor-forming trajectory. Briefly, we filtered our Granger causality analysis to all known genes associated with EMT (EMTdb). This included a list of 1184 genes which served as “seed genes” for the network. Starting with these seed genes, gene dynamics data was used to identify if they acted as upstream drivers of other genes or downstream targets of other TFs within our dataset. To expand the network, we included two degrees of causal connectivity (p < 0.05) around seed genes. Regardless of directionality (upstream or downstream), causality was visualized as causal TF to target interactions. To refine the causal network, we compared it with high-confidence transcriptional interactions from TRUSST v2 (Transcriptional Regulatory Relationships Unraveled by Sentence-based Text mining (https://www.grnpedia.org/trrust/)) database, which was used as the canvas for all known regulatory relationships across the human genome. All networks were generated using Cytoscape 3.9.0 (https://cytoscape.org/).

## Code Availability

The Cflows package, as implemented in python, is available for download with a guided tutorial on the Krishnaswamy Lab Github page: https://github.com/KrishnaswamyLab/cflows.

## Data Availability

The scRNA-seq datasets in this study will be made available for download on the NCBI Gene Expression Omnibus (GEO): https://www.ncbi.nlm.nih.gov/geo/.

## Biological Methods Details

All measurements are taken from distinct samples.

### Cell Culture

MDA-MB-231, HCC1806, HCC38 cells were purchased from ATCC and cultured in DMEM (MDA-MB-231 cells) or RPMI (HCC1806 and HCC38) supplemented with 10% (v/v) FBS (PS; 5.000 units penicillin and 5 mg streptomycin/ml in H_2_O, Sigma Aldrich, cat no. P4333). Cells were routinely tested to confirm the absence of mycoplasma contamination. All cell line-specific media were supplemented with 1% (v/v) penicillin-streptomycin.

### Flow cytometry

Bulk HCC38 cell lines were cultured in 2D tissue culture dishes. For isolating CD44^*hi*^cells from this bulk population, cells were trypsinized and stained with CD44 antibody (BD Biosciences anti-human CD44-PE-cy7 (1:800)) for 25 min at 4C. CD44^*hi*^cells were sorted on BD Aria III. Sorted cells were cultured in media supplemented with 0.1% (v/v) gentamicin and 1% (v/v) antibiotic-antimycotic for at least two passages to avoid contamination. Multiple rounds of FACS enrichment were performed on these expanded cultures until pure CD44^*hi*^populations were isolated (Figure S3a). To identify EPCAM^+/-^, CAV1^+/-^, EPCAM/CAV1^+/+^ populations, HCC38 CD444^*hi*^were subjected to FACS sorting (n = 3) using Anti-CD326 (EPCAM) (1:50, Invitrogen #53-8326-42) and Anti-Caveolin 1 (1:50, BD Biosciences, Figure S3b). For cell cycle experiments, FUCCI-transduced cells were analyzed to identify G1 (RFP+), G1/S transition (RFP+/GFP+), and S/G2/M (GFP+) populations (Figure S3c). Data acquisition was performed using BD Aria III and FACSDiva software (BD Biosciences and data analysis was performed using Flowjo X10.7.1.

### Tumorsphere assay

Single-cell suspensions were plated in ultra-low attachment 96-well plates (Corning^®^ 96-well Clear Flat Bottom Ultra-Low Attachment Microplate #3474) at low densities optimized to ensure tumorspheres arose from single anchor-independent cells. HCC1806, unsorted HCC38 cells, FUCCI labelled, and HCC38 CD44^*hi*^cells were seeded in 100 *µ*L at 100 cells/well. Cell-line specific serum-free media was supplemented with 1% (v/v) penicillin/streptomycin, 20 ng/ml EGF, 20 ng/ml FGFb, 4 *µ*g/mL heparin, 1x B27, and 1% (v/v) methylcellulose (Sigma-Aldrich). Fresh media was topped up every 5 days by adding 50 *µ*L per well of the appropriate tumorsphere media. For the scRNA-seq analysis, HCC38-seeded samples were collected at days 0, and then at days 2, 12, 18, and 30 in TSA culture.

For functional assays, tumorspheres were counted at days 7, 12, or 30 using manual counting of whole wells under brightfield at 4X magnification or using Incucyte^®^ S3, Sartorius (Spheroid mode). Total tumoursphere count per well, when counting manually or Object Count (per mm^2^) when using Incucyte S3 software was used as the metric for quantification. A two-tailed unpaired t-test was used to calculate significance for siControl and siESRRA or vehicle and ESRRAi conditions and was plotted using GraphPad Prism Version 9.3.1.

For 3D immunostaining, tumorspheres were fixed with 10% formalin (Australian Biostain Pty Ltd) for 1 hour at RT in a gently rocking rotator and washed in TBS (3 × 15 min). Tumorspheres were then permeabilized with 100% methanol for 10 minutes at 4°C and washed in TBS (3 × 15 min). Tumorspheres were then blocked in TBS 5% BSA, 10% Horse Serum and 0.1% Triton O/N at 4 degrees with rotation. Following incubation with primary antibodies ESRRA (Cell Signalling Technology, E1G1J, 1:200, Cat no. 13826); ZEB1 (Santa Cruz, #H-102, 1:200); CDH1 (BD Biosciences/(36/E-Cadherin, Cat. no. 610181, 1:200) diluted in blocking buffer O/N at 4°C in rotation, spheres were washed (4 × 30 min) in TBS and stained with the appropriate fluorophore-conjugated secondary antibodies (1:500) (Ms Cy3 (#M30010), Rb 647 (#A32722), Ms 488 (#A11001), Rb 488 (#711546152), ThermoFisher Scientific) and DAPI O/N at 4°C. Tumorspheres were washed with TBS (4 × 30 min) then resuspended in 20 *µ*L of mounting media (ProLong Diamond Antifade Mounting Media (ThermoFisher Scientific)) and mounted between a glass slide and a coverslip spaced by tape.

Labelled tumorspheres were imaged using confocal microscopy (Leica DMI 6000 SP8 with 40x (NA 1.3) or 63x (NA 1.4) oil objectives or a Nikon A1R confocal with 20x Plan Apochromat air objective (NA 0.75) at 2x zoom using an HD25 resonance scanner) using identical acquisition settings (optimised per protein marker) for all time points. Quantitative image analysis was performed using CellProfiler (v4.2.1, [58]) to segment individual cells via maximum cross-entropy-based threshold detection of nuclei (DAPI) and cell bodies (sum of all channels). Mean intensity per cell was measured for each marker, with per cell image and quantitative data integrated via a Knime software (v4.6.4, [59]) for visualisation [60,61].

### ESRRA knockdown and validation assays

Predesigned siRNA specific to ESRRA and scrambled siRNA were purchased from Integrated DNA Technologies, USA (TriFECTa^®^ RNAi Kit, Design ID hs.Ri.ESRRA.13). HCC38 CD44^*hi*^cells were seeded in 24 well plates at a density of 9000 cells/well and ESRRA knockdown was performed using 20nM of pooled siRNA (hs.Ri.ESRRA.13.1, hs.Ri.ESRRA.13.2 and hs.Ri.ESRRA.13.3) using Lipofectamine RNAiMax (ThermoFisher Scientific, USA) as per the manufacturer’s protocol. Cells were harvested for protein extraction 48hr post siRNA transfection to study the downstream effect on CDH1 expression upon ESRRA knockdown using western blot. ESRRA was knocked down in HCC38 CD44^*hi*^, HCC38 CD44^*lo*^, MDA-MB-231 and HCC1806 cells using 20nM of pooled siRNA (ThermoFisher Scientific siRNA ID: s4829 and s4830) using Lipofectamine RNAiMax (ThermoFisher Scientific, USA) as per the manufacturer’s protocol. Cells were harvested after 24hrs of transfection to seed tumorspheres or 48hr after transfection for protein extraction. Cells transfected with 20nM Silencer™ Select Negative Control No. 1 (ThermoFisher Scientific) were used as controls.

In parallel, HCC38 CD44^*hi*^cells were also treated with an ESRRA inhibitor ((2-Aminophenyl)(1-(3-isopropylphenyl)-1H-1,2,3-triazol-4-yl)methanone) (BLD Pharm, China, Cat no. BD01201330) [referred to as Compound 14 (C14)] at 5 and 10*µ*M concentrations. Vehicle controls were treated with DMSO. Cells were harvested 48 hrs post-treatment and processed for protein extraction, which was studied for changes in ESRRA and CDH1 expression using western blot (n=3 biological replicates). Vehicle and C14-treated cells were harvested after 24 hours of transfection to seed tumorspheres.

### Western Blotting

Proteins were extracted from control and treated cells using ice-cold modified RIPA buffer (50 mM Tris-HCl pH 7.5, 150 mM NaCl, 1 mM EDTA, 1 mM EGTA, 1% Triton X-100, 0.1% SDS with supplemented with 1X protease and phosphatase inhibitor cocktails). The lysates were sonicated using a QSONICA Q55 probe sonicator at 50 kHz for 20 seconds in an ice bath. This whole cell lysate was centrifuged at 14,000 x g for 10min at 4°C and stored in −80°C until further use. Proteins were quantified using the Pierce™ BCA Protein Assay Kit (Cat. 23227, Thermo Fisher Scientific, USA) as per the manufacturer’s instructions. 20*µ*g of whole cell lysates were separated on 1D SDS-PAGE using NuPAGE™ 4 to 12%, BisTris (Cat. NP0322BOX, ThermoFisher Scientific, USA). Proteins separated on the gel were transferred onto nitrocellulose membrane using Trans-Turbo Transfer system (BioRad Laboratories, USA) at 1.3V for 7min. The membrane was blocked using 5% milk in TBS 1hr. The membrane was next incubated overnight at 4°C with primary antibodies against the protein of interest [1: 1000 ESRRA (Cell Signalling Technologies, USA, Cat. no. 13826), 1:1000 CDH1 (Cell Signalling Technologies, USA, Cat. no. 3195S), 1:5000 GAPDH (Cell Signalling Technologies, USA, Cat. no. 97166S), 1:5000 *β*-Actin (Cell Signalling Technologies, USA, Cat. no. 3700S)]. After incubation with the primary antibody, membranes were washed with 1X TBST for 3 × 10min on a rocker at room temperature. Next, membranes were incubated with appropriate HRP-conjugated secondary antibodies. Washing steps were repeated, and the membrane was developed using ECL substrate (Western Lightning™ Ultra, Perkin Elmer or Clarity Western ECL Substrate) and scanned using Fusion FX Vilber Lourmat scanner. Each Western blot experiment was performed using three biological replicates, to calculate the statistical significance (p-value) of relative fold-change in expression. GAPDH or *β*-Actin was used to normalize the relative fold-change expression value of ESRRA and CDH1. The signal intensity of the bands in western blots was quantified using Image Studio Lite version 5.2 (LI-COR Biotechnology, USA). Normalised signal intensity readouts were plotted on GraphPad Prism Version 9.3.1. Statistical significance of fold-change protein expression between siControl and siESRRA was calculated using a two-tailed unpaired t-test.

### Animal Studies

All mice were 8-9 weeks of age at time of injections. 4×10^5^ MDA-MB-231 luc cells (siControl or siESRRA) were injected in 100 *µ*L saline via tail vein injections in each mouse (n=12/arm). siRNA transfection was performed as described before using 10nM siControl or 10nM pooled siESRRA (siRNA ID: s4829 and s4830, ThermoFisher Scientific). Colonization of injected cells to lungs was captured through live *in-vivo* bioluminescence imaging (BLI) on Day 0, 2, 7 and 11 using PerkinElmer IVIS Spectrum In Vivo Imaging Systems. Lungs from 5 mice/arm were harvested at Day 2 and were subjected to H&E staining. This resulted in 7 mice/arm for the remaining experiment until Day 11. During each imaging session, a dosage of 10 *µ*L luciferin/g mouse body weight [D-luciferin Potassium Salt stock (7.5 mg/ml, Sigma, LUCK-2G)] was injected subcutaneously. Imaging was performed 5 min after D-luciferin administration animals (auto-exposure and 1sec. C and B magnification) to maintain uniform photon flux. Normalized fold change radiance values for each day was calculated by dividing the raw radiance readings (p/sec/cm^2^/sr) of each animal on a given day by raw radiance value of animal#1 in control on that day. Normalised fold change radiance was plotted on GraphPad Prism Version 9.3.1. An unpaired t-test was performed to calculate statistical significance of normalised fold change radiance values between siControl and siESRRA arms for each day.

For testing if inhibition of ESRRA sensitises TNBC cells to chemotherapy, MDA-MB-231-*luc* cells, (1×10^6^ cells) were prepared in 25 *µ*L 20% Matrigel (BD Matrigel Matrix Growth Factor Reduced (GFR) / serum-free media were injected into the inguinal mammary fat pads of these mice. Twelve (Vehicle control and C14 single arm) to fifteen animals (chemotherapy and combination therapy) were enrolled per treatment arm for each experiment. Animals were randomized into treatment arms at the time tumors ranged between 50-150 mm^3^. Tumor volume was estimated by caliper measurements twice per week up until the harvest date. ESRRA inhibitor C14 was administered by oral gavage (C14, 25 mg/kg) for 4 days a week for 3 weeks in one treatment cycle. C14 was dissolved in DMSO and resuspended in sesame oil at a final concentration of 6.25mg/ml. Mice were administered 100 *µ*L of the drug resuspension per 25g of body weight. Vehicle or chemotherapy single treatment arms were administered diluent by daily oral gavage for the duration of the treatment cycles. Chemotherapy (20 mg/kg Doc) was administered once per week via intraperitoneal injections on day 2 of the 4-day treatment regimen with ESRRA inhibitor/week. Vehicle and chemotherapy single-treatment arms received saline injections on day 2 of each week within the treatment cycle. Animals were harvested when they reached the ethical endpoint (1000 mm^3^ or tumor burden greater than 10% of the animal’s body weight). Growth kinetics of the tumor were monitored through live *in-vivo* bioluminescence imaging (BLI) using PerkinElmer IVIS Spectrum In Vivo Imaging Systems using the protocol described in the previous section. A two-way ANOVA multiple comparison test was used to compare growth kinetics across four different treatment arms. When comparing 2 treatment arms, a paired t-test was used instead. One-way ANOVA and Tukey’s multiple comparison test were applied to determine differences among tumor weights, tumor volume evolution in response to treatment. A paired t-test was used to compare differences in flux intensity between DTX and DTX + C14 treatments at the end of the treatment cycle.

All experiments were approved by conducted following the National Health and Medical Research Council Statement on Animal Experimentation, the requirements of New South Wales State Government legislation, and the rules for animal experimentation of the Biological Testing Facility of the Garvan Institute and the Victor Chang Cardiac Research Institute (protocol #18/12).

## Supplemental Information

**Table S1:** Left and Middle: comparison of RMSE between ground truth and inferred trajectories in PHATE space and the gene space (lower is better). PHATE Space RMSE is scaled by 10^−3^. Right: comparison of EMD between ground truth and predicted held-out populations (lower is better). EMD values are scaled by 10^−2^.

| Method↓ | Data→ | PHATE Space RMSE |  |  | Gene Space RMSE |  |  | Held-Out Population EMD |  |  |
| --- | --- | --- | --- | --- | --- | --- | --- | --- | --- | --- |
|  |  | Bifurcation | Cycle | Linear | Bifurcation | Cycle | Linear | Bifurcation | Cycle | Linear |
| **BEMIOflow (ours)** |  | **2.60** | **0.777** | **3.40** | **0.0773** | **0.118** | **0.0853** | **0.465** | **0.465** | **0.935** |
| TrajectoryNet [3] |  | 6.16 | 2.49 | 8.08 | 0.150 | 0.151 | 0.118 | 1.07 | 0.710 | 1.50 |
| OT-CFM [15] |  | 2.98 | 3.79 | 5.33 | 0.0842 | 0.209 | 0.0960 | 0.495 | 0.818 | 1.32 |
| SB-CFM [15] |  | 2.83 | 1.58 | 5.67 | 0.0835 | 0.141 | 0.0978 | 0.516 | 0.526 | 1.28 |

**Table S2:** Mean and standard deviation of the graph edit distance between the inferred graph and the ground truth, calculated over *n* = 5 sets of trajectories inferred using BEMIOflow (lower is better).

| Method | Gene Regulatory Network (SERGIO) |  |  |
| --- | --- | --- | --- |
|  | Bifurcation | Cycle | Unidirectional |
| **GC (ours) [9]** | **35.3 ± 1.7** | **75.2 ± 2.3** | **120.8 ± 3.4** |
| OCE [16] | 65.6 ± 4.2 | 130.9 ± 6.3 | 190.3 ± 6.9 |
| PC [17, 18] | 72.4 ± 4.1 | 145.7 ± 7.2 | 200.2 ± 7.6 |
| mTE [18] | 48.6 ± 3.3 | 110.4 ± 4.1 | 145.3 ± 4.6 |
| mMI [19] | 42.2 ± 2.7 | 90.6 ± 3.2 | 130.1 ± 4.7 |
| NRI [20] | 140.3 ± 15.6 | 255.4 ± 25.8 | 310.7 ± 22.3 |
| DCRNN [21] | 155.8 ± 20.5 | 285.6 ± 30.2 | 395.1 ± 35.7 |
| GTS [22] | 165.4 ± 22.3 | 300.8 ± 33.4 | 440.2 ± 40.9 |
| NIR [23] | 160.6 ± 18.4 | 270.3 ± 28.5 | 390.5 ± 32.8 |

**Supplementary Figure S1:**
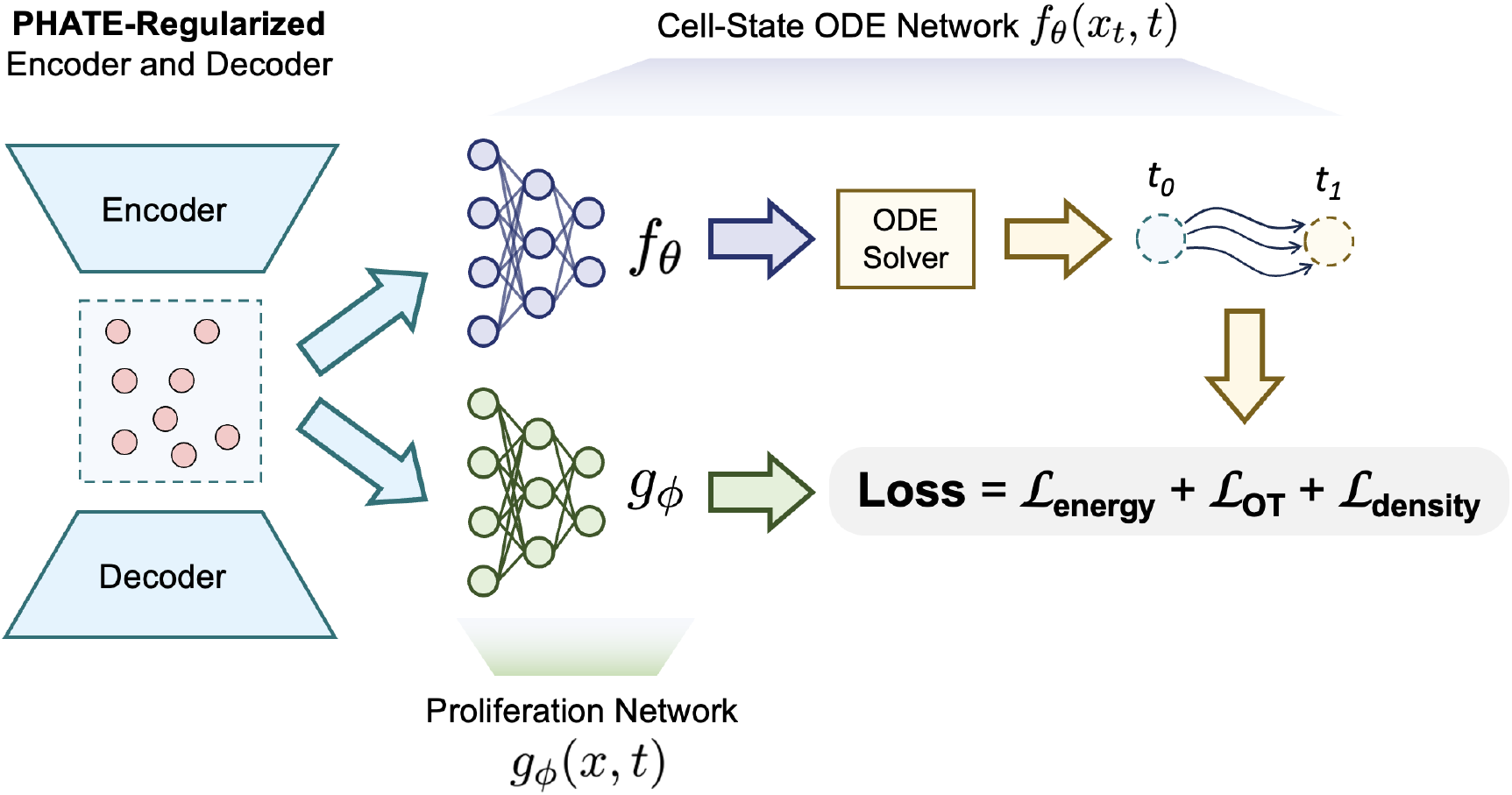
Schematic of BEMIOflow. BEMIOflow simultaneously models the cell trajectories using a cell-state ODE network *f*_*θ*_(*x*_*t*_, *t*) and the growth rate using a proliferation neural network *g*_*ϕ*_(*x, t*). These modeling are performed in the latent space of the PHATE-regularized encoder and decoder. The loss function is a combination of an energy term, an optimal transport term, and a density term. More details can be found in the Methods section.

**Supplementary Figure S2:**
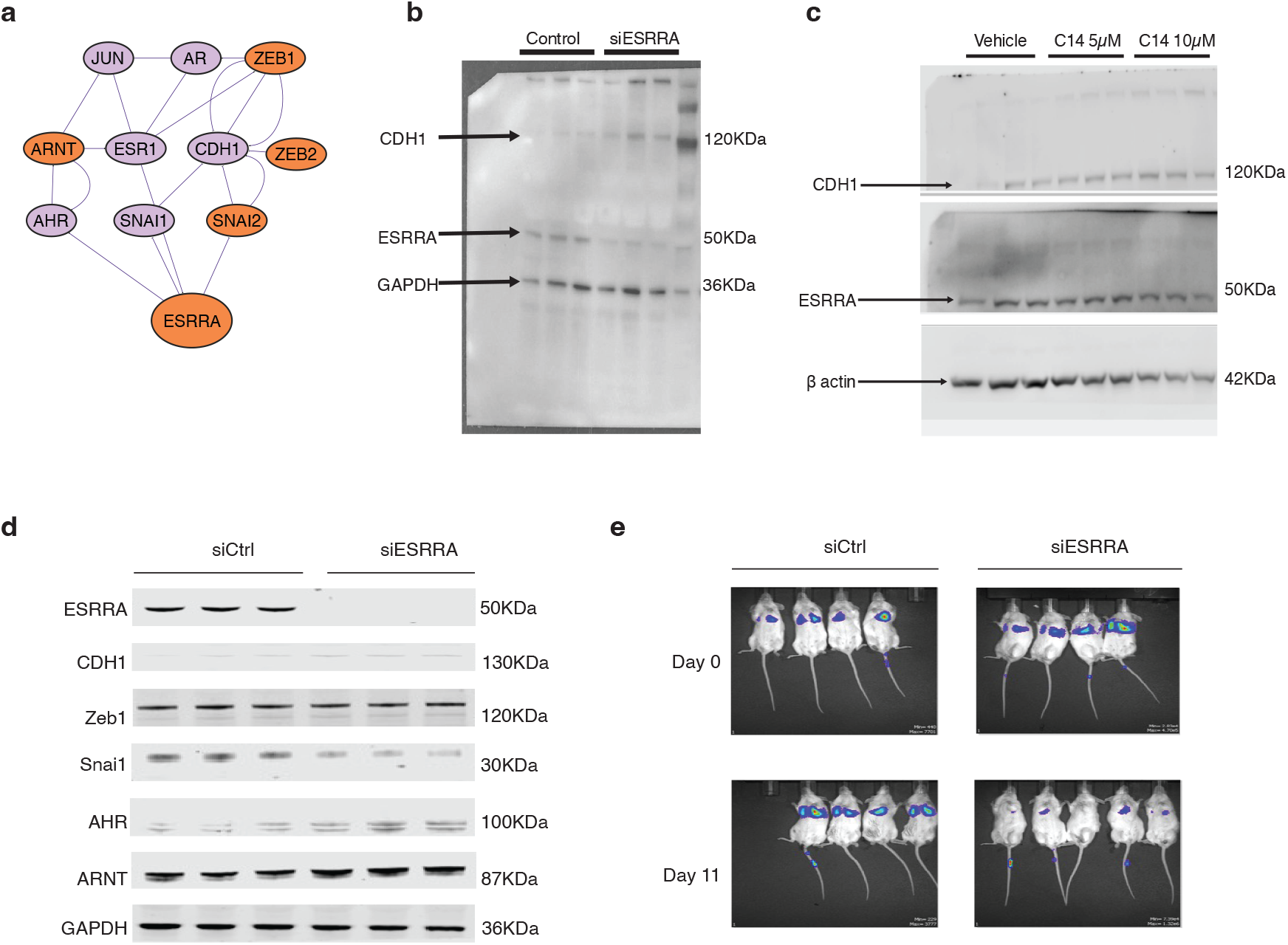
ESRRA regulates a subnetwork of genes critical for the emergence of the epithelial cell state. **a**, ESRRA regulates a subnetwork of genes, including known EMT transcription factors such as SNAI1/2, ZEB1/2, and CDH1. **b**, Western blot of siCtrl and siESRRA whole cell lysates from HCC38 CD44hi cell line showing downregulation of ESRRA corresponds to upregulation of CDH1 **c**, HCC38 CD44hi cells treated with ESRRAi (C14) at 5 and 10*µ*M concentration show upregulation of CDH1 in comparison to vehicle-treated cells. **d**, Western blot of siCtrl and siESRRA whole cell lysates from MDA-MB-231 cell line showing protein expression of genes regulated within the ESRRA subnetwork **e**, Representative bioluminescence images of mice injected with siCtrl and siESRRA MDA-MB-231-luc cells at Day 0 and Day 11 after injection.

**Supplementary Figure S3:**
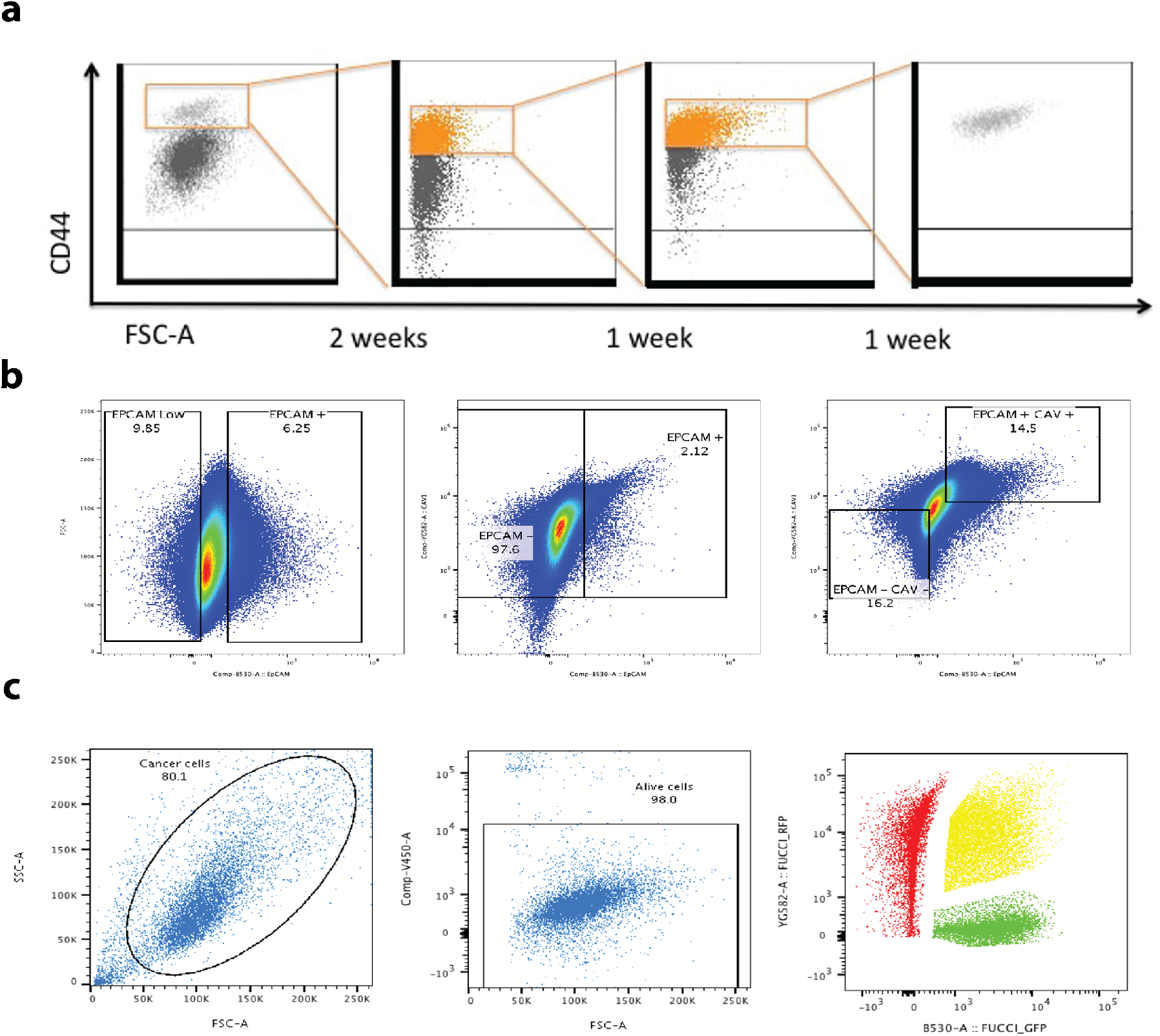
Gating strategies for the isolation and analysis of HCC38 cancer cell subpopulations. **a**, Gating strategy for the enrichment and isolation of the CD44^*hi*^ subpopulation from bulk HCC38 cells. The gate on the highest CD44-expressing cells (orange box) was refined over multiple rounds of fluorescence-activated cell sorting (FACS) to achieve a pure population. **b**, Strategy for the isolation of EPCAM^+^/CAV1^+^ double-positive cells from the CD44^*hi*^ line. Sequential gates were applied to select for cells expressing both the Epithelial Cell Adhesion Molecule (EPCAM) and Caveolin-1 (CAV1) surface markers. **c**, Gating strategy for cell cycle analysis using the FUCCI (Fluorescent Ubiquitination-based Cell Cycle Indicator) system. Cells were first gated to exclude debris (FSC vs. SSC) and dead cells (viability dye). Live, FUCCI-transduced cells were then separated into G1 (RFP^+^), G1/Stransition (RFP^+^/GFP^+^), and S/G2/M (GFP^+^) phases based on fluorescence.

## Notes

### Competing Interest Statement

The authors have declared no competing interest.

### Summary of Updates

The computational analysis has been revised with an updated tool called MIOflow in place of an older tool called Trajectorynet.

